# Human Cardiac Organoids to Model COVID-19 Cytokine Storm Induced Cardiac Injuries

**DOI:** 10.1101/2022.01.31.478497

**Authors:** Dimitrios C Arhontoulis, Charles Kerr, Dylan Richards, Kelsey Tjen, Nathaniel Hyams, Jefferey A. Jones, Kristine Deleon-Pennell, Donald Menick, Diana Lindner, Dirk Westermann, Ying Mei

**Author notes:** **Corresponding Authors**, Ying Mei, PhD.

## Abstract

Acute cardiac injuries occur in 20-25% of hospitalized COVID-19 patients. Despite urgent needs, there is a lack of 3D organotypic models of COVID-19 hearts for mechanistic studies and drug testing. Herein, we demonstrate that human cardiac organoids (hCOs) are a viable platform to model the cardiac injuries caused by COVID-19 hyperinflammation. As IL-1βis an upstream cytokine and a core COVID-19 signature cytokine, it was used to stimulate hCOs to induce the release of a milieu of proinflammatory cytokines that mirror the profile of COVID-19 cytokine storm. The IL-1 β treated hCOs recapitulated transcriptomic, structural, and functional signatures of COVID-19 hearts. The comparison of IL-1β treated hCOs with cardiac tissue from COVID-19 autopsies illustrated the critical roles of hyper-inflammation in COVID-19 cardiac insults and indicated the cardioprotective effects of endothelium. The IL-1β treated hCOs also provide a viable model to assess the efficacy and potential side effects of immunomodulatory drugs, as well as the reversibility of COVID-19 cardiac injuries at baseline and simulated exercise conditions.

## Introduction

Besides pulmonary complications, one of the most prevalent complications of COVID-19 is acute cardiac injury (ACI) ^1,2^. 20% to 25% of hospitalized COVID-19 patients have suffered ACI, which are associated with a poor prognosis and increased mortality rate ^3,4^. Despite the evidence of direct viral infection of the myocardium ^5,6^, a growing body of literature suggests COVID-19 induced cytokine storm is a major contributor to the ACI ^7-10^. Unlike the cytokine storm induced by Chimeric Antigen Receptor T (CART) Cell therapy ^11,12^, the inflammatory profile of COVID-19 cytokine storm is marked by lymphopenia, granulocytosis, and a milieu of proinflammatory cytokines with a strong innate component. Several innate cytokines such as IL-1β, and IL-6 have been used clinically as markers of patient prognosis and offer viable options for pharmacologic intervention ^13-15^.

Despite the critical roles of COVID-19 cytokine storm in ACI, the current lack of animal and *in vitro* models has limited the mechanistic understanding and impeded drug development. COVID-19 has prompted testing of previously approved immunomodulatory drugs in clinical trials ^16–24^. Given the difficulty in obtaining human heart biopsies, these trials are limited by their inability to reveal molecular responses of human hearts to drug treatments. Additionally, clinical data has suggested COVID-19 survivors with ACI can experience long-term cardiac abnormalities ^25,26^. As follow-up studies may take years to complete, this highlights an unmet need to develop effective models that can predict long-term cardiac outcomes of convalescent COVID-19 patients to provide guidance for clinical monitoring and therapeutic interventions ^27^.

Recently, our laboratory has developed *in vitro* 3D human organoids (hCOs) that are composed of human pluripotent stem cell-derived cardiomyocytes (hPSC-CMs), human cardiac fibroblasts (hcFBs), human umbilical vein endothelial cells (HUVECs), and human adipose derived stem cells (hADSCs) ^28^. These hCOs provide a powerful *in vitro* system to model cardiovascular pathologies. For example, they have been shown to recapitulate the transcriptomic, structural and functional hallmarks of myocardial infarction and cardiovascular disease exacerbated pharmacotoxicity ^29^.

IL-1 β is one of the first cytokines released in response to viral infection by macrophages and epithelial cells ^12^. IL-1 β serum levels have strongly correlated with severe COVID-19 disease, despite a short half-life in serum ^13^. Lung and heart tissues of COVID-19 patients have revealed increased expression of the IL-1 β receptor (IL1R1), corroborating high levels of IL-1 β signaling ^30^. As an upstream cytokine, IL-1 β is known to induce the release of downstream cytokines including IL-6 ^12,31^. Given all of the cell-types comprising the hCOs (e.g., HUVECs, hPSC-CMs, hcFBs) can produce cytokines in response to proinflammatory stimulation ^32-35^, we postulated that IL-1 β stimulation would induce a cytokine storm in our organoids, recapitulating the hyperinflammation affecting COVID-19 hearts and inducing functional and morphological changes consistent with clinical findings. As IL-1 β may be considered a nonspecific upstream stimulus, we used clinical data and autopsy samples to validate the experimental design of using IL-1 β treated hCOs to model COVID-19 specific cardiac pathologies. Herein, we report that treating hCOs with 1 ng/mL IL-1 β induces the release of an array of proinflammatory cytokines that mirrors the profile of COVID-19 cytokine storm. IL-1 β treated hCOs recapitulated transcriptomic, structural, and functional hallmarks of the COVID-19 induced acute cardiac injuries. We further illustrate that IL-1 β treated hCOs provided a valid model to test the effects of immunomodulatory drugs to treat cardiac injuries caused by COVID-19 cytokine storm. Lastly, we showed that IL-1β treatment induced cardiac dysfunction is reversible if given sufficient recovery time, indicating the possible recovery of COVID-19 induced cardiac injuries.

## Results

### RNA Seq Analysis of IL-1β Treated hCOs Established their Transcriptomic Relevance with COVID-19 hearts

This study was designed on the growing premise that systemic inflammation appears to be the major driving force leading to COVID-19 ACI (**Fig. 1**) ^7–9^. As IL-1β is an upstream proinflammatory cytokine and a signature core COVID-19 cytokine, we treated hCOs with IL-1β (1 ng/mL) to simulate the COVID-19 ACIs using the treatment regimen depicted in **Fig. 2A**. Four days was selected for IL-1 β treatment because clinical data has indicated that it takes 12-96 hours to transition from moderate to severe disease characterized by hypercytokinemia ^36^. We did not observe significant differences in hCO diameter and gross morphology after IL-1β treatment (**Fig. 2B**). There was no significant difference between normalized TUNEL positivity between control and IL-1 β treated hCOs (**Fig. 2C**), consistent with minimal evidence of cardiomyocyte death in COVID-19 autopsies ^37^.

**Figure 1.**
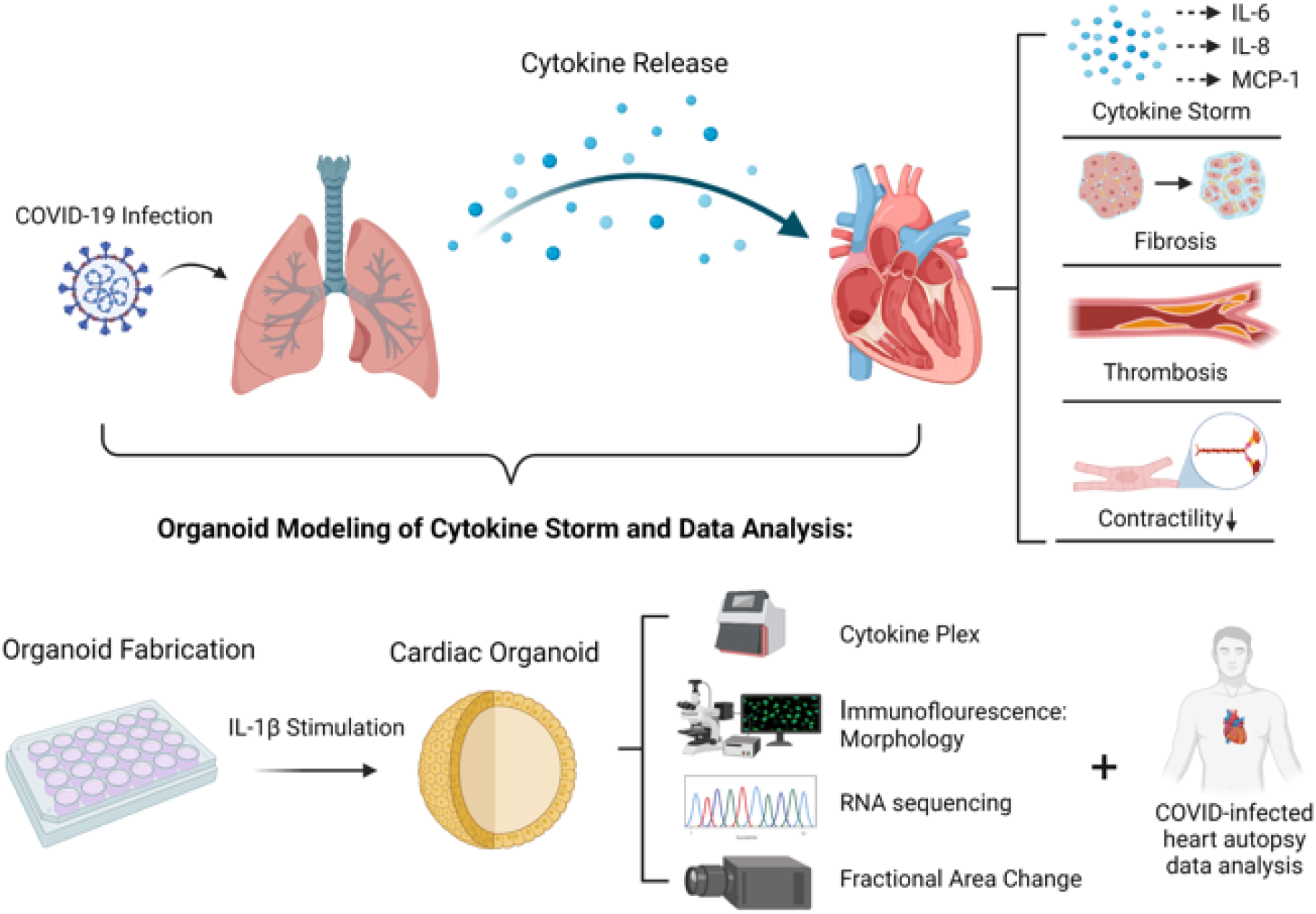
Premise of Physiology and Experimental Design. SARS-CoV2 infects patient’s lungs, inducing cytokine production and release into the blood stream. Cytokines that have entered the blood stream affect the heart and induce fibrosis, thrombi, and reductions in contractile function. Human Cardiac Organoids (hCOs) stimulated with IL-1β were multiplexed for downstream cytokine release, alterations in morphology and transcriptomic signatures, and assessed for reductions in contraction amplitude. Transcriptomic signatures of hCOs were also compared to COVID-19 autopsy samples.

**Figure 2.**
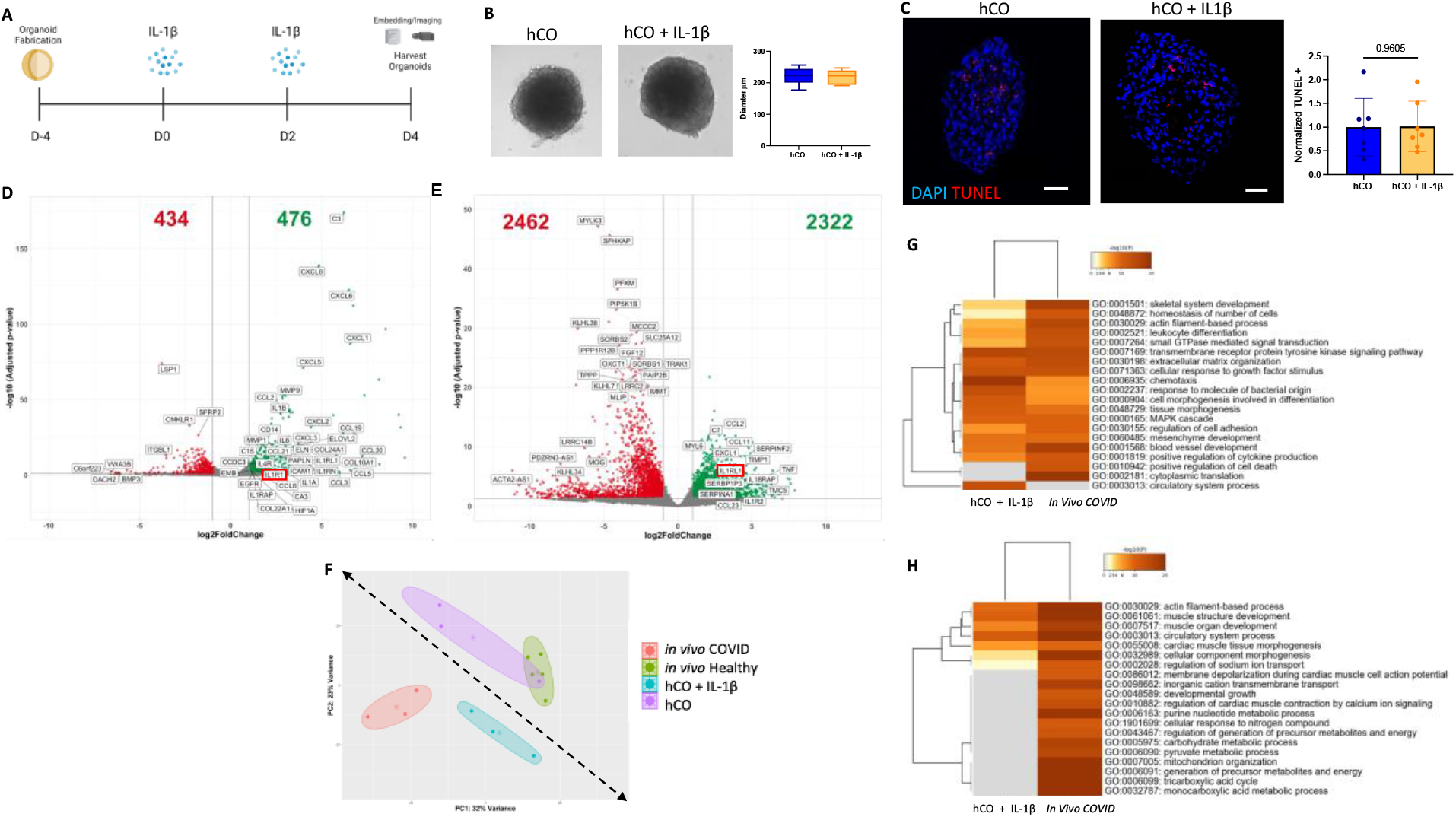
Transcriptomic Signature of Human Cardiac Organoids Treated with IL-1β Resemble that of COVID-19 Autopsy Samples. (A). Schematic of treatment regimen for human cardiac organoids. (B) (Left) Representative brightfield images of Organoids on D4 (Right) Diameters of organoids on Day 4 (mean +/− s.d.); n = 10, p = 0.9290 by student’s t-test (C) Representative images of immunofluorescent staining of human hCOs and hCOs + IL-1β. (Blue = DAPI, Red = TUNEL). Scale bar = 50 um for each image. Quantification of Normalized TUNEL expression to the right (mean +/− s.d.); n =6, p = 0.9605 by student’s t-test. (D) Volcano plot illustrating DEGs between hCOs and IL-1β treated hCO. (E) Volcano plot illustrating differentially expressed genes (DEGs) of publicly available dataset from COVID-19 autopsy samples, relative to control patients. (F) Principal Component Analysis (PCA) of showing clustering of hCOs and control patients, and COVID-19 patients with treated hCOs + IL-1β. (G) Pathway analysis comparing upregulated pathways in IL-1β treated human cardiac organoids and autopsy samples of COVID-19 patients. (H) Pathway analysis comparing downregulated pathways in pathways in IL-1β treated human cardiac organoids and autopsy samples of COVID-19 patients.

To establish system level relevance of IL-1 β treated hCOs for modeling COVID-19 hearts, we performed RNA sequencing (RNAseq) on hCOs with/without IL-1 β treatment and compared their transcriptomic profiles to COVID-19 and healthy heart autopsy samples from publicly available datasets ^38^. We first plotted the top differentially expressed genes (DEGs) for our IL-1β treated hCOs (compared to control hCOs) and the COVID-19 hearts (compared to healthy controls). As seen in **Fig. 2D-E**, IL1R1 was upregulated in both IL-1β treated hCOs and COVID-19 hearts, supporting the critical role of the IL-1 β signaling in the COVID-19 induced hyperinflammation. Principal Component Analyses (PCA) showed control hCOs and healthy hearts grouped in the top right, while the IL-1 β treated hCOs and COVID-19 hearts grouped together in the bottom left in the PC1/PC2 plot (**Fig. 2F**), illustrating the high fidelity of IL-1β treated hCOs to model COVID-19 hearts. This is further supported by the heatmap of the shared top 20 unregulated and downregulated pathways between IL-1 β treated hCOs (vs. control hCOs) and COVID-19 hearts (vs. healthy hearts). IL-1β treatment of hCOs led to the upregulation of several key pathways that were also upregulated in the COVID-19 hearts (**Fig. 2G)**. The top Gene Ontology (GO) terms included leukocyte differentiation (GO:0002521), chemotaxis (GO:0006935), and positive regulation of cytokine production (GO:0001819), indicating a similar proinflammatory transcriptomic profile between IL-1 β treated hCOs and COVID-19 hearts. Moreover, the shared extracellular matrix organization and mesenchyme development indicated the IL-1β treated hCOs recapitulated cardiac fibrosis observed in COVID-19 patients ^39^. In **Fig. 2H**, hCOs and COVID-19 hearts also shared common downregulated genes, which were aligned with key GO terms such as “muscle organ development” (GO:0007517), and “cardiac muscle tissue morphogenesis” (GO:0055008). The downregulated cardiomyocyte structure pathways are consistent with acute reductions in cardiac output (acute heart failure, cardiogenic shock in COVID-19 patients with acute cardiac injuries ^40^. A significant portion of differences in the downregulated pathways between the IL-1β treated hCOs and COVID-19 hearts is associated with metabolism (e.g., tricarboxylic acid cycle, mitochondrion organization), which was attributed to underdeveloped mitochondria of immature hPSC-CMs compared to adult human cardiomyocytes ^41^. In summary, the RNAseq analyses established similar transcriptomic profiles between IL-1 β treated hCOs and COVID-19 hearts.

### IL-1β treated hCOs Recapitulated Cytokine Profile from Severe COVID-19 Patient Serum

We next examined whether the IL-1 β treated hCOs were capable of secreting key proinflammatory cytokines found in the serum of severe COVID-19 patients. In particular, IL-6 is a pro-inflammatory cytokine thought to be a key marker of cytokine storm. Its levels have strongly correlated with both disease severity and mortality of COVID-19 patients ^13,14,42–44^. To confirm IL-6 was released in response to IL-1β treatment, supernatant was collected on day 4 after IL-1β treatment and IL-6 levels were significantly upregulated, with a mean fold change of 6.5 (**Supplementary Fig. 1A-B**), consistent with the measurements from the serum of severe COVID-19 patients (6-8-fold) and distinct from that of CART cytokine storm (~75 fold) ^11^. To assess the direct effects of IL-6 on our human cardiac organoids, IL-6 +/− its soluble receptor (sRα) were added to our cardiac organoids for 10 days at 10 ng/mL, 50 ng/mL and 100 ng/mL (**Supplementary Fig. 1C**). No significant changes in contraction amplitude were found at any dose, with or without the addition of its soluble receptor, suggesting that in the context of the COVID-19 hearts, IL-6 may serve as a marker rather than the etiology for cardiac dysfunction. Given the increase of IL-6, supernatants from IL-1 β treated hCOs collected were examined to determine the induced cytokine profile present using a bead-based multiplexed immunoassay system (Eve Tech, Canada). GM-CSF, IL-1β, IL-2, IL-4, IL-5, IL-6, IL-8, and MCP-1 were all upregulated (**Fig. 3A**), consistent with previous reports detailing cytokine composition in the COVID-19 cytokine storm ^14,15,45^. Though TNFα, IFNγ, and IL-13 were measured, they were not as consistently upregulated in our samples (**Supplementary Fig. 2**).

**Figure 3.**
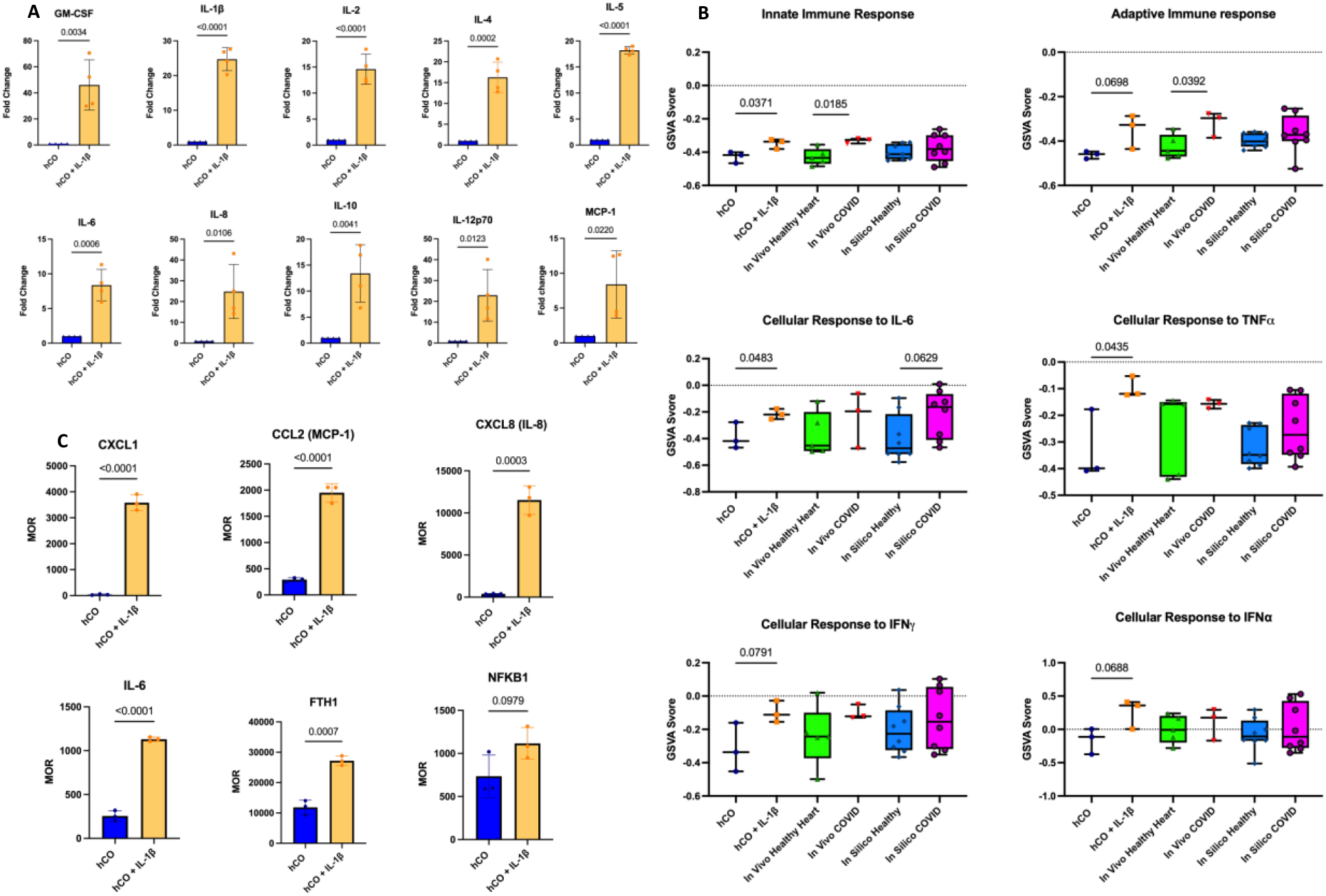
Inflammatory Profile of IL-1β treated Human Cardiac Organoids Resembles that of COVID-19 Cytokine Storm. (A) Cytokine multiplex results showing fold change of cytokines in D4 supernatant as compared to hCOs; (mean +/− s.d.), n =4, compared using student’s t-test. (B) GSVA analysis of hCOs and IL-1β treated hCOs (n= 3), Healthy *In Silico* and COVID *In Silico* (n = 5, and n = 3, respectively) and samples from Healthy and COVID autopsy samples (n = 8) assessing multiple GO terms pertaining to inflammation (mean +/− s.d.). Analysis performed using student’s t-test. (C) Median of Ratios (MOR) of key genes upregulated upon IL-1β treatment; n =3 (mean +/− s.d.), analysis performed by student’s t-test.

To support the multiplexing analyses, we evaluated the RNAseq data of the IL-1 β treated hCOs and COVID-19 hearts on inflammatory GO terms and pathways using Gene Set Variation Analysis (GSVA) analyses. To account for the cellular composition differences between cardiac tissues and hCOs (e.g., without immune and neural cells), we utilized publicly available single cell RNAseq (scRNAseq) data from COVID-19 and healthy hearts to develop *in silico* hCOs ^30,46^. The *in silico* hCOs were constructed by selecting the specific cell types (i.e., ventricular cardiomyocytes, ventricular fibroblasts, cardiac endothelial cells, and cardiac pericytes) and compositions similar to those used in hCOs fabrication and aggregating them into an *in silico* organoid-like pseudo-bulk sample per donor (**Supplementary Fig. 3)**. hCOs showed significant increases in GSVA score upon IL-1β treatment under GO terms of “Innate Immune Response” (GO:0045087), “Cellular Response to IL-6” (GO:0071354), and “Cellular Response to TNF” (GO:0071356) (**Fig. 3B**), mirroring the clinical findings of the upregulation of innate immunity, IL-6 and TNFα signaling observed in COVID-19 hearts ^15,42,47^. Though not significant, IL-1β treatment promoted the increase of a variety of proinflammatory pathways in hCOs such as “Adaptive Immune Response” (GO:0002250), “Cellular Response to IFNα” (GO:0035457) and “Cellular Response to IFNγ” (GO:0071346), with a similar trend observed in the *in silico* COVID hCOs and COVID-19 heart samples. Importantly, IL-1 β treatment led to the elevation of key proinflammatory innate genes such as CXCL1, CCL2, and CXCL8 in the hCOs. IL-1 β treatment also upregulated key clinical markers of inflammation such as IL-6 and FTH1 (**Fig. 3C**). NF_κ_B1, a key transcriptional regulator of cytokine production and other key biological processes showed increasing trends with IL-1β treatment. Collectively, these immune-related genes support the GO terms and pathway analyses.

### IL-1β treated hCOs showed reduced cardiac function and pathological cardiac fibrosis

Given the similarities of the IL-1β treated hCOs to *in vivo* samples (i.e., *in silico* hCOs and heart samples) under immune-related Go terms, we next performed GSVA analyses for both cardiac structure and function related GO terms (**Fig. 4A**). We observed a decrease with IL-1β treated hCOs of key cardiac structural and functional GO terms such as “Cardiac Myofibril Assembly” (GO:0055003) and “Cardiac Muscle Contraction” (GO:0060048), mirroring the *in silico* COVID-19 organoids and COVID-19 heart samples. To support the results of RNAseq analyses, we examined the effects of IL-1β treatment on the cardiac structure and function of hCOs. As seen in **Fig. 4**, IL-1 β treated hCOs had distinct morphologies (**Fig. 4B**), significantly reduced fractional area change (FAC) (i.e., contraction amplitude) (**Fig. 4C**) and reduced sarcomere width (**Fig. 4D**), consistent with the downregulated cardiac structure and function GO terms. This was also seen at higher doses of IL-1β (**Supplementary Fig. 4A-C**). This was further supported by the significant downregulation of actinin α2 (ACTN2) and β-myosin heavy chain (MYH7), while cardiac troponin I3 (TNNI3) and tropomyosin 1 (TPM1) trended downwards (**Fig. 4E**).

**Figure 4.**
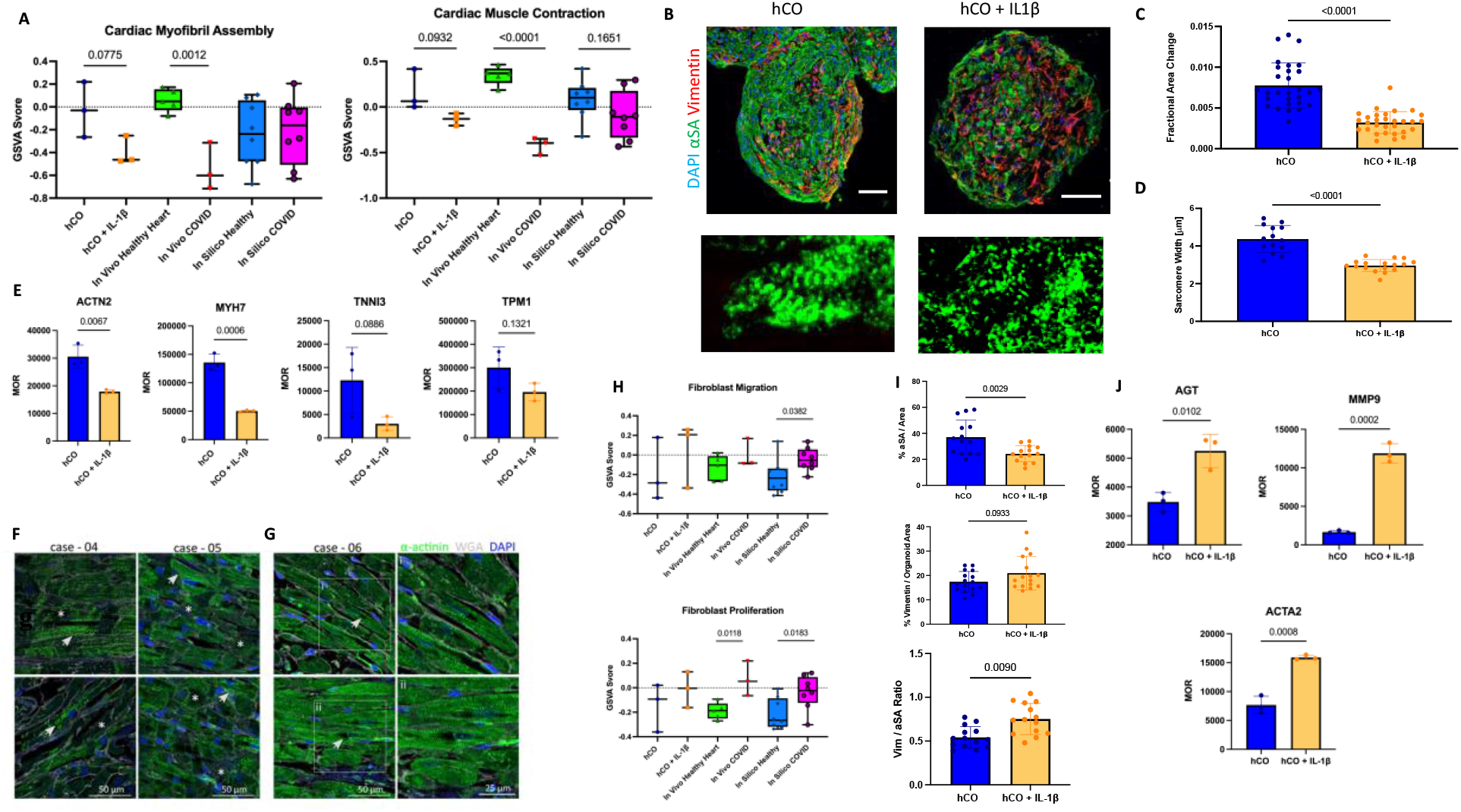
Human Cardiac Organoids Recapitulate Key Hallmarks of Cardiac Function and Fibrotic Response to COVID-19. (A) GSVA analysis of vehicle and IL-1β treated organoids (n= 3), Healthy *In Silico* and COVID *In Silico* (n = 5, and n = 3, respectively) and samples from Healthy and COVID autopsy samples (n = 8) assessing multiple GO terms pertaining to cardiac function, and structure (mean +/− s.d.), analysis performed using student’s t-test. (B) Immunofluorescent staining of either hCOs (top left) or IL-1β treated hCOs (top right) on Day 4 (Green = α-SA, Red = Vimentin, Blue = DAPI). Scale = 50 um for each image. Higher magnification of sarcomere structure from organoids shown for hCOs (bottom left), or IL-1β treated hCOs (bottom right). (C) Fractional Area Change (FAC) shown for either hCOs or IL-1β treated hCOs, (mean +/− s.d., n = 29, 30, respectively) p <0.0001 by student’s t-test. (D) Sarcomere width quantification for either in hCOs or IL-1β treated hCOs (mean +/− s.d., n = 14, and 17, respectively) p < 0.0001 by student’s t-test. (E) key genes involved in cardiac function, Median of Ratios (mean +/− s.d., n = 3 for each group). Student’s t-test performed for each comparison. (F) Two representative α-actinin staining (green) of cardiac tissue section from case-04 and case-05, both without cardiac SARS-CoV-2 infection, are shown. The normal sarcomeric structure of cardiomyocytes is indicated by arrows, whereas inconsistent actinin staining as a sign of sarcomeric disarray is indicated by asterisks. Nuclei counterstained with DAPI are shown in blue, extracellular matrix stained with WGA is shown in white. (G) Two representative images of cardiac tissue from case-06 (without cardiac SARS-CoV-2 infection). White boxes indicted zoomed in areas of the image on the right. Some cardiomyocytes lack nuclear DNA staining (DAPI). White arrows denote the putative region of nuclear localization. (H) GSVA analysis of vehicle and IL-1β treated organoids (n= 3), Healthy *In Silico* and COVID *In Silico* (n = 5, and n = 3, respectively) and samples from Healthy and COVID autopsy samples (n = 8) assessing multiple go terms pertaining to fibrosis and fibroblast behavior (mean +/− s.d), analysis performed by student’s t-test. (I) (Top) Quantification of α-SA staining in organoids. α-SA area divided by organoid area to yield percentage of a-SA per organoid, (mean +/− s.d., n = 16 for both groups, p = 0.0029 by student’s t-test (Middle) Quantification of Vimentin staining in organoids. Vimentin area divided by organoid area to yield percentage of Vimentin per organoid, (mean +/− s.d., n = 16 for both groups, p = 0.0933 by student’s t-test. (Bottom). Ratio of Vimentin area to a-SA area per organoid. (n = 13, and 14, respectively), p = 0.0090 by students t-test. (J) Key genes involved in fibrotic response in the human heart, Median of Ratios (mean +/− s.d., n = 3 for each group). Student’s t-test performed for each comparison.

To assess the validity of the cardiac findings within our hCO model, cardiac tissue was harvested from 95 autopsies of COVID-19 patients. 49 of these patients had confirmed SARS-CoV-2 infection in their cardiac tissue, with viral localization limited to interstitial spaces. 46 of these 95 autopsies had no detectable SARS-CoV-2 in the myocardium. Both SARS-CoV-2 negative (**Fig. 4F-G**), and positive (**Supplemental Fig. 5A-B**) hearts showed decreased α-sarcomeric actinin (α-SA) expression with increased cardiomyocyte destruction, indicating that COVID-19 induced cardiomyocyte damage is independent of viral infection of the myocardium.

Cardiac fibrosis has been reported in autopsies of COVID-19 patients ^39^. The *in silico* COVID-19 hCOs and COVID-19 heart samples showed significant increases in “Fibroblast Proliferation” (GO:0048144) scores with IL-1β treated hCOs showing increases yet high variation (**Fig. 4H**). Notably, the immunofluorescent staining revealed that IL-1 β treated hCOs showed significantly less α-SA expression, increased vimentin expression (p = 0.0933) and a significantly higher vimentin to α-SA ratio (**Fig. 4I**), supporting the cardiac fibrosis. The significant increase in vimentin to α-SA ratio was also seen at higher doses of IL-1β **(Supplemental Fig. 4B**). This is further supported by the upregulation of cardiac fibrosis hallmark genes such as angiotensinogen (AGT), matrix metalloproteinase 9 (MMP9), and α-smooth muscle actin (ACTA2), in the hCOs (**Fig. 4J**).

### IL-1β treated hCOs showed prothrombotic vasculature

Severe COVID-19 patients can experience thrombotic events and vascular damage ^48–50^. To assess the vascular content of our hCOs, we stained our organoids for Platelet Endothelial Cell Adhesion Molecule (PECAM/CD31) and von Willebrand Factor (vWF). The CD31 expression between vehicle and IL-1β treated was not significantly different, while the expression of vWF, a clotting factor, was significantly higher in the IL-1β treated organoids (**Fig. 5A**). Notably, the endothelial cell content was largely depleted in our hCOs at higher IL-1β doses (**Supplemental Fig. 6A-B**), which validated our IL-1 β dose (1 ng/mL) used in the study. While transcriptomic analysis only showed weak indications of a coagulative transcriptome (GO:0050817) of the hCOs upon IL-1 β treatment (**Fig. 5B**), the upregulation of key prothrombotic genes such as Intercellular Adhesion Molecule 1 (ICAM1), E-Selectin, and Vascular Adhesion Protein (VAP1) was significant (**Fig. 5C**), similar to COVID-19 patients ^48^. Additionally, downregulation of Claudin-5 (CLDN5) may lead to vascular permeability ^48^. Nitric Oxide Synthase 3 (NOS3) expression trended downwards in our IL-1 β treated hCOs and indicates a potential decrease in the ability of endothelial cells to support CM function through nitric oxide ^51^. PLAU, a gene responsible for counteracting coagulation, was found to be significantly upregulated, and high levels of its signaling has been found to predict disease severity in COVID-19 patients ^52^.

**Figure 5.**
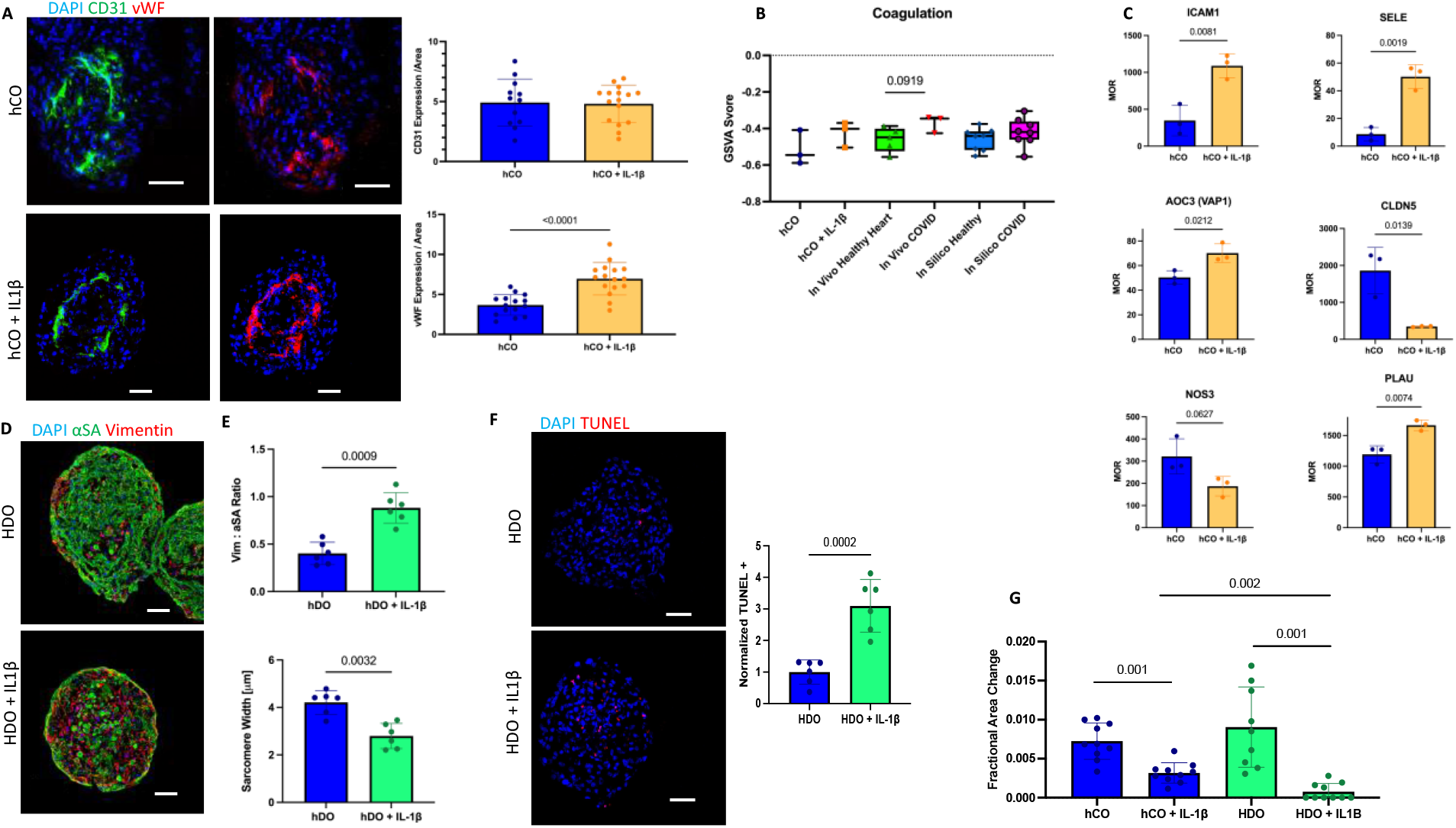
Human Cardiac Organoids Treated with IL-1β Recapitulate Hallmarks of Vascular Response to COVID-19. (A) Immunofluorescent staining of either hCOs (top row) or IL-1β treated hCOs (bottom row) (Green = CD31, Red = vWF). Scale bar for all images is 50 um. Quantification of staining (Top) reveals no significant difference in CD31 content between vehicle and IL-1β treated organoids, (mean +/− s.d., n = 12, and n = 16 p = 0.9969 by student’s t-test. (Bottom) reveals significant increase in vWF expression per organoid in IL-1β treated hCOs by student’s t-test, p <0.0001 (mean +/− s.d., n = 14, and n = 16, respectively). (B) GSVA analysis of hCOs and IL-1β treated hCOs (n= 3), Healthy *In Silico* and COVID *In Silico* (n = 5, and n = 3, respectively) and samples from Healthy and COVID autopsy samples (n = 8) assessing go term pertaining to coagulation (mean +/− s.d.), analysis by student’s t-test. (C) Key genes involved in the vascular response to inflammation/COVID-19 in the human heart, Median of Ratios (mean +/− s.d, n = 3 for each group). Student’s t-test performed for each comparison. (D) Representative Images of immunofluorescent staining of HDOs (left) or IL-1β treated HDOs (right) (Green = α-SA, Red = Vimentin, Blue = DAPI). Scale bar = 50 um for each image. (E) Quantification of Vimentin to α-SA ratio per organoid left, and sarcomere width in um (right). mean +/− s.d., n= 6 for each group. p = 0.0009 and 0.0032 by student’s t-test for Vimentin: a-SA ratio and sarcomere width, respectively. (F) Immunofluorescent staining for TUNEL of either HDO (top) or HDO + IL-1β (bottom), (Red = TUNEL, Blue = DAPI). Scale bar = 50 um for each image. (Right) Quantification of TUNEL expression normalized to the HDO control, (mean +/− s.d., n = 6, p = 0.0002 by student’s t-test.). (G) FAC of either hCOs or HDO organoid formulations treated +/− IL-1β on Day 4. Comparison of contraction amplitude between formulations reveals significant difference (p = 0.002) in FAC between IL-1β treated formulations via student’s t-test (n = 9-10).

Recent studies have indicated endothelial cells are the main target of SARS-CoV2 viral infection in the myocardium, which then induce secretion of proinflammatory cytokines (e.g., IL-1β)^53^. To assess how the endothelial cells affect IL-1β mediated damage to our hCOs, we removed HUVECs from our standard hCO formulation to prepare HUVEC Deficient Organoids (HDOs). Representative images of HDOs and HDOs treated with IL-1β (**Fig. 5D**) showed increases in Vimentin to α-SA ratio and decreases in sarcomere width (**Fig. 5E**), consistent with IL-1β treated HCOs. Unlike the hCOs (**Fig. 2C**), TUNEL expression was significantly increased upon IL-1β treatment in the HDOs **(Fig. 5F)**, indicating a shift in mechanism in IL-1β mediated damage to the organoids. Further, **Fig. 5G** indicates the HDOs had a functional reduction in contraction amplitude that was significantly more severe than that of the hCOs. Collectively, these experiments indicate the critical roles of endothelial cells in mitigating COVID-19 cytokine storm mediated damage.

### IL-1β treated hCOs are a viable platform to test immunomodulatory drugs

We leveraged our hCOs to examine the ability of clinically available immunomodulatory drugs to alleviate the IL-1 β induced cardiac injuries to predict their clinical performance. We chose to assess 4 common immunomodulatory drugs (**Fig. 6A**): an IL-1 receptor antagonist (IL-1RA) (similar mechanism of action as Anakinra); Tocilizumab, a monoclonal antibody against the IL-6 receptor; Baricitinib, a JAK/STAT inhibitor aimed at blocking cytokine receptors; and Dexamethasone, a potent glucocorticoid meant to inhibit the transcription of cytokines, now used commonly to treat hospitalized COVID-19 patients ^54–56^. Organoids were treated with each drug concurrently with IL-1 β on Day 0, and drug was replenished in the media with every media change / cytokine replenishment. As seen in **Fig. 6B**, Dexamethasone was the only drug that was able to ameliorate the hCOs from the IL-1β induced reduction in organoid contractility (i.e., fractional area change). Tocilizumab was unable to improve contraction amplitude, consistent with its modest therapeutic benefits in clinic ^57^. Baricitinib was also unable to improve contraction amplitude, which was also reported by Mills et al when tested in their hCO system ^58^. While effective, Dexamethasone exacerbated the ratio of vWF/CD31, showing even higher levels of vWF than IL-1β treatment alone (p = 0.0514) (**Fig. 6C**). This is consistent with the prothrombotic side effects of Dexamethasone observed in the clinical studies ^59^. (Representative images of CD31/vWF staining for each condition can be found in **Supplementary Fig. 7A-B**). Interestingly, although they were unable to spare the hCOs from IL-1 β induced decreases in contraction amplitude, Tocilizumab and Baricitinib helped preserve sarcomere width (**Fig. 6D**). Moreover, Dexamethasone showed a higher vimentin expression in the hCOs (**Fig. 6E**), indicating an increased presence of fibroblasts. While initially unexpected, literature has suggested that Dexamethasone can induce fibroblast proliferation in certain instances ^60^. In addition to validating the therapeutic benefits of various immunomodulatory drugs, these studies also revealed the potential side effects of the Dexamethasone that deserve clinical monitoring.

**Figure 6.**
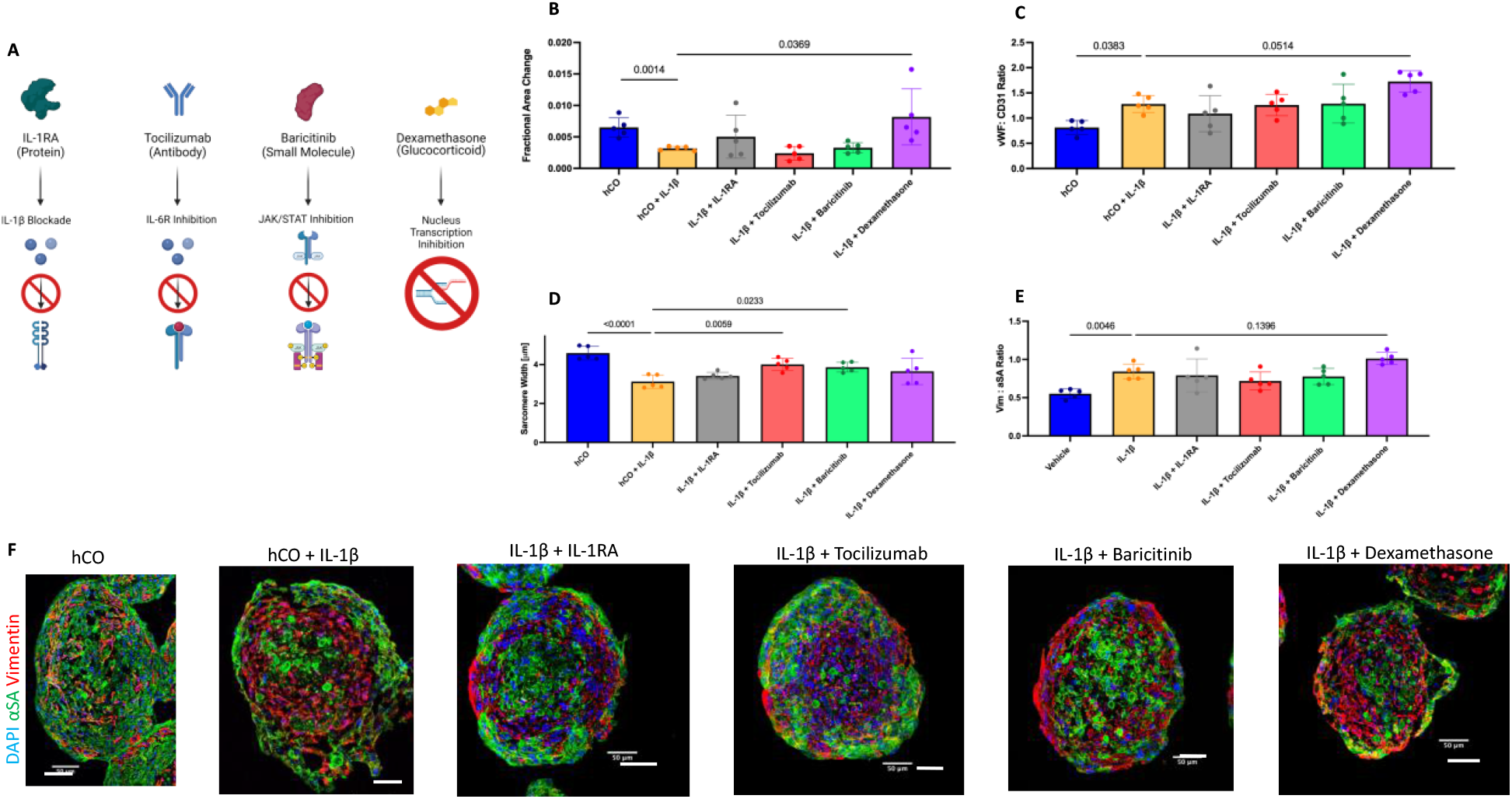
Human Cardiac Organoids Provide Platform for Immunomodulatory Drug Testing. (A) Schematic indicating the mechanism of action for each immunomodulatory drug tested with the human cardiac organoid system. (B) FAC on Day 4 for hCOs, IL-1β treated hCOs, or IL-1β treated hCOs with each immunomodulatory drug. (n = 5 for each group, mean +/− s.d.), analysis performed by student’s t-test. (C) vWF to CD31 ratio for hCOs, IL-1β treated hCOs, or IL-1β treated hCOs with each immunomodulatory drug, (n = 5 for each group, mean +/− s.d.), analysis performed by ANOVA. (D) Sarcomere width in um for hCOs, IL-1β treated hCOs, or IL-1β treated hCOs with each immunomodulatory drug, (n = 5 for each group, mean +/− s.d.), analysis performed by ANOVA. (E) Vimentin to α-SA ratio for vehicle, IL-1β, or IL-1β with each immunomodulatory drug, (n = 5 for each group, mean +/− s.d.,), analysis performed by ANOVA. (F). Representative Images of immunofluorescent staining for hCOs, IL-1β treated hCOs, or IL-1β treated hCOs with each immunomodulatory drug. (Green = α-SA, Red = Vimentin, Blue = DAPI). Scale bar for each image = 50 um.

**Figure 7.**
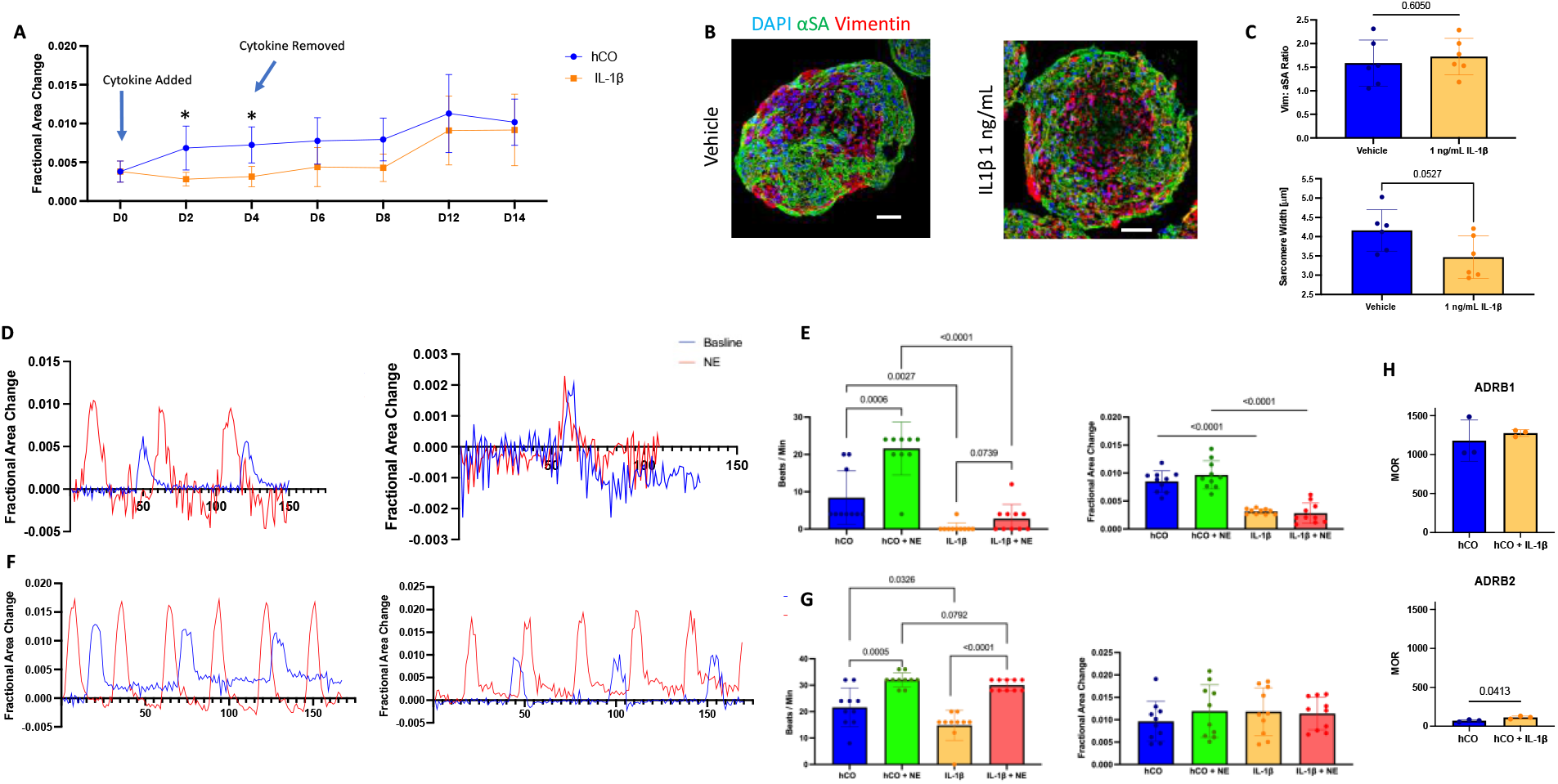
Human Cardiac Organoids Are a Viable Model to Test Recovery Studies. (A) FAC over time, with arrows indicating the addition and removal of cytokine (mean +/− s.d., n = 9-10, * = p<0.05 by students t-test). (B) Immunofluorescent staining of organoids on D14 treated initially with either vehicle or IL-1β (Green = α-SA, Red = Vimentin, Blue = DAPI, Scale bar for each image = 50 um). (C) (Top) Vimentin to α-SA ratio per organoid for D14 organoids (Bottom) Quantification of sarcomere width for D14 organoids. (n = 6 for each group, p values = 0.6050 and 0.0527, respectively by student’s t-test. (D). Waveform of fractional area change with Day 4 hCOs (left) or IL-1β treated hCOS (right) in the presence (red) or absence (blue) of 1 uM Norepinephrine. (E) (Left) Number of beats per minute of hCOs, hCOs stimulated with NE, IL-1β treated hCOs, or IL-1β treated hCOs stimulated with NE. (Right) FAC for each group before and after NE stimulation. (n = 10 for each group, mean +/− s.d., and analyses performed by student’s t-test.) (F) Waveform of fractional area change with Day 14 hCOs (left) or IL-1β treated hCOs (right) in the presence (red) or absence (blue) of 1 uM Norepinephrine. (G) (Left) Number of beats per minute in hCOs, hCOs treated with NE, IL-1β treated hCOs, or IL-1β treated hCOs stimulated with NE. (Right) FAC for each group before and after NE stimulation. (n = 10 for each group, mean +/− s.d., and analyses performed by student’s t-test.) (H) Median of Ratios of the beta 1 (top) and beta 2 (bottom) adrenergic receptors in hCOs and IL-1β treated hCOs, (mean +/− s.d., n =3), analysis performed by student’s t-test.

### IL-1β treated hCOs provide a valid platform for recovery studies

Significant interests surround the reversibility of COVID-19 ACIs and the extent to which patients may be susceptible to long-COVID as they recover from active infection and enter convalescence ^61^. To investigate COVID-19 ACI reversibility, hCOs were conditioned with a recovery period (normal culture medium with no IL-1β) for 10 days after 4 days of IL-1β treatment (**Fig. 7A**). It took 2 days of recovery before FAC was not statistically significant, despite their mean FAC values not leveling out until day 14. Representative images for the control hCOs and the hCOs recovered from IL-1β treatment are shown in **Fig. 7B**. Despite similar mean FACs and vimentin/α-SA ratios for each treatment on day 14, the recovered hCOs after IL-1β treatment had a lower mean sarcomere width (**Fig. 7C**) and reduced CD31 and vWF staining **(Supplemental Fig. 8**).

To assess the functional recovery of hCOs on D14, we performed a “Stress Test” to simulate physiological exercise. While hCOs responded to norepinephrine (NE) stimulation by showing a significant increase in spontaneous beating rate, and IL-1 β treated hCOs did not respond to NE stimulation on Day 4 (**Fig. 7D-E**). However, both groups responded to the NE stimulation on day 14 (**Fig. 7F-G**), with each group showing a significant increase in the beating rate. Of note, levels of the adrenergic receptor β1were not different via RNAseq analysis between vehicle and IL-1 β, though levels of the β2 receptor were significantly increased **(Fig. 7H**). These studies indicated the cardiac injuries caused by the COVID-19 cytokine storm could be reversed after a sufficient recovery period.

## Discussion

The cytokine profile seen from the IL-1 β treated hCOs suggests the organ level response to IL-1 β closely mirrors that of the serum cytokine profile frequently reported in literature. Based on these findings, we shed light on the possibility that the inflammatory response of the myocardium either mirrors the systemic inflammatory profile measured via serum analysis, or that the myocardium itself contributes to the systemic inflammation found in severe COVID-19 patients as indicated in recent publications ^62^. Mills et. al. recently published a study using a cytokine screen on hCOs to identify specific cytokine induced diastolic dysfunction ^58^. They ultimately used a cocktail of IL-1β, IFNγ, and poly(I:C) as their “cytokine storm” formulation and assessed effects of TNFα on hCOs. Our data may indicate that a multi-cellular response to IL-1β shares sufficient downstream mediators to capture hallmarks of both TNFα and IFNγ, despite the lack of a consistent upregulation of either gene. Of note, IL-1 β treated organoids showed a marked upregulation of Urokinase Type Plasminogen Activator. Its receptor, Plasminogen Activator Urokinase Receptor (PLAUR), has been found to indicate increased risk of progression of COVID-19 to respiratory failure. A recent phase 3 clinical trial indicated success with Anakinra when using these soluble Urokinase Plasminogen Activator Receptor (suPAR) levels to guide treatment, highlighting the key role of IL-1 signaling in the development of COVID-19 severe disease ^52^.

Cardiac fibrosis following COVID-19 has been reported, even in mild or asymptomatic patients, which may lead to cardiac complications such as arrhythmias later in life ^61,63,64^. As long-term effects of COVID-19 continue to be assessed in patients, understanding the reversibility of cardiac dysfunction has become paramount, particularly in the context of acute convalescence. Currently, clinical guidelines recommend patients recovering from COVID-19 infection should not resume physical activity for at least 10 days after the onset of symptoms and 7 days after symptom resolution ^65^. Our data indicates it takes up to 2.5 times of the exposure time for contractile machinery to be re-upregulated. Even at the point of functional recovery, sarcomere width was still reduced, supporting the clinical guidelines. Though the reversibility of these injuries may be increased due to the use of hPSC-CMs, reversibility of IL-1β mediated cardiac dysfunction has been shown both *in vitro* and *in vivo^51,66,67^*.

While recent studies suggested that SARS-CoV2 viral infection of endothelium has led to the IL-1β production and myocardial dysfunction ^62^, the results from the HDO system indicates the endothelium also mitigates the COVID-19 cardiac insults, consistent with its well-established cardioprotective effects^68^. Interestingly, recent COVID-19 autopsy studies have shown direct SARS-CoV2 virial infection of myocardium is associated with significant reduction of endothelial cells and early death ^53^. As current COVID-19 clinical trials that target endothelium have been focusing on anti-coagulation therapies ^69^, our data indicates the importance of endothelium protection to ameliorate the COVID-19 induced cardiac injuries.

In summary, our results showed IL-1 β induced the release of a milieu of the proinflammatory cytokines from hCOs, with a similar profile to COVID-19 cytokine storm. Our data also validated the IL-1β treated hCOs’ ability recapitulate the hallmarks of the transcriptome, structure, and function of COVID-19 hearts. We further demonstrated the IL-1 β treated hCOs are an effective testing platform for immunomodulatory drugs and long-term reversibility of COVID-19 induced cardiac pathologies. Given the validated response to IL-1 β, our study opens the possibility for patient-specific cells to be incorporated into our hCOs to help identify the genetic variations that contribute to the large spectrum of COVID-19 responses observed.

A limitation of our system is the lack of immune cells. To address this, we introduced the concept of *in silico* hCOs by leveraging the publicly available scRNAseq data of COVID-19 hearts and well-defined cell type and composition of our hCOs (**Supplementary Fig. 3**). The IL-1 β treated hCOs showed higher similarity with COVID-19 hearts than the *in silico* COVID-19 organoids in multiple GSVA analyses (**Fig. 3B, 4A, 4H, and 5B**). While this can be attributed to patient variation and the use of bulk RNAseq (used on the hCOs and *in vivo* heart samples) rather than scRNAseq (used to construct the *in silico* hCOs), our findings indicate the significance of the immunomodulatory properties (e.g., cytokine release) of cardiomyocytes, endothelial cells, and stromal cells in the organoids and in COVID-19 hearts. Additionally, because we utilized hPSC-CMs, it is possible the hCOs can better withstand IL-1 β mediated damage. However, we noticed similar trends in cardiac contractile structure and function upon exposure to IL-1 β, demonstrating that hPSC-CMs recapitulate the key phenotypic shifts seen in adult human cardiomyocytes. Though our findings are consistent with hPSC models of direct SARS-CoV-2 infection ^6^, direct comparisons are needed to validate our findings.

## Methods

### Cell Culture

hPSC-CMs (iCell Cardiomyocytes, Cellular Dynamics) were cultured according to the manufacturer’s protocol. iCell Cardiomyocytes (donor 01434) were used for all experiments. Briefly, hPSC-derived CMs were plated on a 0.1% gelatin coated 6 well plates in iCell Cardiomyocyte Plating Medium (CDI) at a density of 3-4 × 10^5^ cells per well and incubated at 37 °C in 5% CO_2_ for 4 days. 2 days after plating, the plating medium was replaced with iCell Cardiomyocyte maintenance medium (CDI). After 4 days of monolayer preculture, cardiomyocytes were lifted using trypLE Express (Gibco Life Technologies) and prepared for organoid fabrication. Human cardiac ventricular fibroblasts (CC-2904, 0000401462, Lonza) were cultured in FGM-2 medium (Lonza), at passages 3-4 for organoid fabrication. Human umbilical vein endothelial cells (HUVECS; C2519A) were cultured in EGM-3 medium (Lonza) and were used passages 2-3 for organoids fabrication. Human adipose-derived stem cells (hADSCs; PT-5006, 0000410257, Lonza) were cultured in low-glucose Dulbecco’s modified Eagle’s medium supplemented with 10% FBS and 1% Penicillin-streptomycin, 1% glutamine and 1% antimycin (Gibco Life Technologies). hADSCs were used at passages 3-5 for organoid fabrication.

### Fabrication of Organoids

We have previously described the fabrication of our organoids ^28,70^. Briefly, agarose molds fabricated from commercial master micromolds from Microtissues were used as molds for microtissue fabrication, with each mold containing a 7 × 5 matrix of recesses. Organoid cellular suspensions are composed of 55% hPSC–CMs, 24% hcFBs, 14% HUVECs, and 7% hADSCs in medium at a concentration of 2 × 10^6^ cells per mL. To generate organoids with a diameter ~150 μm, 75 μL of the organoid suspension was added into the molds and allowed to settle for 15 minutes. Upon settling, 2 mL of medium was added to submerge the molds in a 12-well plate. Media was changed every 2 days for the entirety of the experiment. The organoids were allowed to form for 4 days, when it was denoted as D0 of the experiment. IL-1 β treatment protocol was then initiated for 4 days. Organoid media is composed of a ratiometric combination of cellspecific medium reflecting the starting cell ratio of the organoid. The hPSC-CM-specific component was defined as glucose-containing F12/DEME medium with 10% FBS, 1% glutamine and 1% non-essential amino acids (Gibco). In the instance of the HUVEC Deficient Organoids (HDOs) the 14% of the HUVEC cell content was replaced with cardiomyocytes in its cell suspension.

### Contraction Analysis

Videos of spontaneously beating organoids from each group were recorded 10 minutes after removing the 12 well plate from the incubator. This was done to equilibrate plates with room temperature and reduce variation in temperature mediated changes in beating. Recording of the videos was performed with a Carl Zeiss Axiovert a1 Inverted Microscope and Zen 2011 software (Zeiss). Threshold edge-detecting in ImageJ software (US National Institutes of Health NIH)) was used on high contrast organoid picture series and graphed over the number of frames. Each beating profile was used to calculate the fractional area change amplitude (fractional change in area of the organoid at maximum contraction and relaxation.)

### Bulk RNA Sequencing

RNA-seq was performed on hCOs using previously described methods^29,71^. Briefly, total RNA was isolated from 35 hCOs per replicate (i.e., one agarose mold per replicate), 18 days after fabrication using the Omega bio-tek E.Z.N.A Total RNA kit (Omega bio-tek, Inc.; Norcross, GA, USA; Cat# R6834-01) with the addition of the Omega homogenizer columns (cat# HCR003). 125 ng of total RNA was used for the construction of the library using the New England Biolabs NEBNext^®^ Poly(A) mRNA Magnetic Isolation Module (Cat# 7490L) and Ultra II Directional RNA Library Prep Kit for Illumina (Cat# 7760L) according to the manufacturer’s instructions. Indexed libraries were pooled and sequenced at VANTAGE (Vanderbilt University Medical Center) on an Illumina NovaSeq 6000(Illumina, Inc.; San Diego, CA, USA), producing paired-end 150 bp reads at a sequencing depth of 25 million reads/sample. Paired-end reads were aligned to the hg38 human reference genome (Genecode GRCh38.p13) using RNA STAR^72^ (v2.7.8a) and subsequent gene counts were generated using htseq-count (v13.5) utilizing the Galaxy Project online platform (v21.01, https://galaxyproject.org/).

### Differential Gene Expression

The R package, DESeq2 (v1.24.0) [doi: 10.1186/s13059-014-0550-8], was used to perform the differential gene expression analysis. Human *in vivo* myocardium samples containing healthy myocardium (n = 5) and myocardium from COVID infected patients^*38*^ (n = 3) were retrieved on the National Institute of Health’s Gene Expression Omnibus (GEO), GSE169241. Differential gene expression analysis was performed using Rstudio (v1.3.1093) (R language v3.6.1). The following two analyses were performed using DESeq2: (1) IL-1β treated hCOs compared to control hCOs and (2) COVID infected patient myocardium compared to healthy myocardium patients. DEG’s that exhibited a log2fold change greater than or less than 1 and an adjusted *p*-value < 0.05 were considered in our study. Volcano plots for each study were generated using the ggplot2 package (v3.3.5)^73^. For pathway analysis, differentially expression genes from both DE analyses were loaded into Metascape^74^ to conduct a multiple gene list analysis for enrichment of Gene Ontology biological processes. For individual gene analysis, analyses were assessed using DESeq2 normalized values (median of ratios) and subsequent t-tests were performed.

Gene expression counts of our COVID-19 cardiac datasets (i.e., hCO, *in silico* and *in vivo)* were batch corrected using ComBat_seq^75^ in the Surrogate Variable Analysis R package(v3.35.2)^76^ and subsequently normalized to log2(counts per million) using the R package “EdgeR”(v3.34). Gene ontology gene sets for GSVA were retrieved from Molecular Signatures Database (v7.4)^77,78^. GSVA was performed using the “gsva” package (v1.40.1) and t-tests were performed on GSVA enriched values. Hierarchical clustering of GO terms were performed using the heatmap package (v1.0.12).

### *In Silico* Organoids

Single nucleus cell RNA-sequencing data from healthy^46^ and COVID hearts^30^ were used for *in silico* organoid fabrication to make pseudo-bulk RNAseq samples per donor. For each donor, ventricular cardiomyocytes (CMs), ventricular fibroblasts (FBs), cardiac endothelial cells (ECs), and pericytes were chosen as representative cell types and maximally randomly sampled to reach the human cardiac organoid ratios of 55% CMs, 24% FBs, 14% ECs, and 7% pericytes. For example, if Donor A had 3000 CMs, 1000 FBs, 1000 ECs, and 1000 pericytes, then 2291 randomly sampled CMs, 1000 FBs, 583 ECs, and 292 pericytes were used for Donor A *in silico* organoid construction. Pericytes were used to represent ADSCs from the hCOs in this case due to ADSC’s pericyte-like characteristics in vivo and in hCOs^79,80,28^. The gene counts were then summed across all cells per donor to obtain pseudo-bulk samples. Counts per million (cpm) normalization was applied, and genes with less than 1 cpm across 70% of the samples were filtered out. This resulted in ~15,000 genes per covid silico organoid and ~23,000 genes per healthy heart *in silico* organoid.

### Fluorescence Imaging and analysis

Organoids were harvested and flash frozen in Tissue-Tek Optimal Cutting Temperature (OCT) compound (Sakura). Embedded organoids were cryosection (7 μm thick) onto glass slides for immunofluorescence staining. Section fixation took place in precooled acetone (−20° C for 12 minutes. After two washes in Phosphate Buffer Saline (PBS) with 0.1% Triton X-100 (Sigma) (PBST), blocking buffer was made with 10% serum corresponding to the host species of the secondary antibodies in PBST. Blocking buffer was added to sections for 1 hour at room temperature. The sections were then washed 2 times (5 minutes) with PBST and stained for primary antibody at 1:200 in PBST for 1 hour at room temperature: mouse anti-alpha sarcomeric actinin (ab9465, Lot: GR3174517-4, Abcam), mouse anti-alpha smooth muscle actin, (A6228, Lot: 056M4828V, Sigma), mouse anti - CD31 (PECAM-1) (cat: 550389, Lot: 5170510, BD Biosciences), rabbit anti-vimentin (ab92547, Lot: GR3258719-11, Abcam), rabbit anti – von Willebrand factor (ab6994, Lot: GR3180938-1, Abcam). Sections were then washed 2 times (5 minutes) with PBST and stained with either complementary secondary antibody or conjugated primary antibodies diluted in PBST (1:200) for 1 hour at room temperature: Alexa Fluor 488 phalloidin (cat: A12379, Lot: 1871076, Invitrogen), goat anti-mouse Alexa Fluor 647 (ab150115, GR3297002-1, Abcam, goat anti-rabbit Alexa Fluor 488 (ab150077, Lot: GR3313703-1, Abcam). After washing with PBST 2 times (5 minutes), nuclei were stained with NucBlue (R37606, Lot: 2165134, Invitrogen) diluted in PBST for 20 minutes at room temperature. Sections were then washed 3 times (3 minutes) in PBST, and coverslips were added using Fluoroshield (Lot: MKCG4762, Sigma) and stored at 4°C until imaging. A TCS SP5 AOBS laser-scanning confocal microscope (Leica Microsystems) was used to image microtissue sections for which z stacks with a thickness of 3-4 μm and a step size 1 μm was used. Intensities for alpha-sarcomeric actinin, vimentin, CD31, and vWF were calculated using ImageJ Threshold Detection after splitting images into individual channels. Threshold area was then measured and normalized to the cross-sectional area of each organoid being measured in the image. To measure co-localization of F-actin and α-smooth muscle actin, a Color Threshold was applied to each image using ImageJ to account for the superimposition of each stain (yellow to measure for green and red overlap). The color threshold area was then measured for each organoid. Sarcomere width was measured by using the line tool in ImageJ. Each individual replicate is the mean of 10 sarcomere widths from one organoid. The Roche In Situ Cell Death Detection Kit (Lot:41569800, Sigma) was used to visualize apoptotic cells in frozen sections of cardiac organoids on the basis of the manufacturer’s protocol. Briefly, sections were fixed with 4% paraformaldehyde in PBS for 20 minutes at room temperature. Sections were washed for PBS for 30 minutes at room temperature, the sections were then incubated with a permeabilization solution (PBS with 0.1 Triton X-100 and 0.1% sodium citrate) for 2 minutes on ice. 50 μL of the TUNEL solution (90% label, 10% enzyme) was then applied to the sections, and incubated for 1 hour at 37°C. Sections were then washed twice with PBS for 5 minutes. Nuclei were then stained with NucBlue (R37606, Lot: 2165134, Invitrogen) diluted in PBS for 20 minutes at room temperature. Sections were then washed 3 times (3 minutes) in PBS, and coverslips were added using Fluoro-Shield (Sigma) and stored at 4°C until imaging. A TCS SP5 AOBS laser-scanning confocal microscope (Leica Microsystems) was used to image microtissue sections for which z stacks with a thickness of 3-4 μm and a step size 1 μm was used. ImageJ particle analysis was used to count the number of DAPI events and TUNEL positive events, from which viability and percent apoptotic cells could be calculated.

### Cytokine Measurements

Supernatants were collected from organoid culture on Day 4. Human Interleukin 6 concentrations were obtained via ELISA (Invitrogen, EH2IL6). ELISA was performed according to the manufacturer’s protocol. Concentrations are reported as mean +/− standard deviation.

### Eve Tech Cytokine Plexing

Supernatants were collected from organoid culture on day 4 and were analyzed for cytokine levels using a Human Cytokine Array Proinflammatory Focused 13 – plex (Eve Technologies, Calgary, AB).

### Cytokine Treatment

IL-1β (Cat:4128-10, Lot: P1098, Biovision) was added to the organoid culture on days 0 and 2 upon media replacement at either 1 ng per mL, 50 ng per mL or 100 ng per mL. IL-6 (Cat 4143-20, Lot:4D17L41430, Biovision) and its soluble receptor, IL-6 sRα (Cat 7102-10, Lot:3H11L71020, Biovision) or both were added on days 0, 2, 4, 6, and 8 upon media replacement. Each cytokine was reconstituted at the recommended stock concentration (with recommended solvents) concentration per the manufacturer’s instructions.

### Drug Testing

Drugs were added to organoids concurrently with IL-1 β on Days 2 and 4 upon media replacement: Tocilizumab (Cat: A1447-200, Lot: 6G02A14470, Biovision), Human IL-1RA (Cat:4263-100, Lot: 8L024263, Biovision), and Dexamethasone (Cat: D1961, Lot: DMGTH-QG, Tokyo Chemical Industry (TCI)) were added to the culture at a concentration of 1 μg per mL. Baricitinib (Cat: 2842-10, Lot: 6H13L28420, Biovision) was added to the culture at 1μM. Each drug was reconstituted at the recommended stock concentration (with recommended solvents) per the manufacturer’s instruction.

### Stimulation with Norepinephrine

Organoids were allowed to equilibrate to room temperature for 10 minutes prior to video recording (as previously described in Contraction Analysis Methods). Individual organoids were filmed using a Carl Zeiss Axiovert a1 Inverted Microscope and Zen 2011 software (Zeiss) for 15 seconds. The number of complete contractions was recorded for each condition. Contraction analysis was then performed. 1 uM of Norepinephrine was then added to the organoids at room temperature and allowed to incubate for 20 minutes. The number of beats within the 15 second were recorded again. Contraction analysis was also performed post NE stimulation.

### Human cardiac tissue specimens

Deceased with confirmed SARS-CoV-2 infection were autopsied at the Institute of Legal Medicine at the University Medical Centre Hamburg-Eppendorf in Germany between April and May 2020^53^. Confirmation of SARS-CoV-2 infection was proven prior to death or post-mortem by quantitative reverse transcription-polymerase chain reaction from pharyngeal swabs. For subsequent analysis, two tissue specimens were collected from the free left ventricular wall during autopsy and either snap frozen in liquid nitrogen or fixed in 10% neutral-buffered formalin. Cardiac SARS-CoV-2 infection was determined as described previously^5^.

### Immunofluorescence staining of cardiac tissue

For immunofluorescence staining, left ventricular tissue was dehydrated and embedded in paraffin. Subsequently, four micometer-thick formalin-fixed paraffin embedded sections were prepared, deparaffinised and rehydrated. A heat-induced antigen retrieval was performed with citrate buffer (pH 6) and sections were permeabilized in 0.2% Triton X-100/tris-buffered saline (TBS) for 10 min. To quench autofluorescence, 0.25% Sudanblack / 70% ethanol was used. Next, sections were blocked with 3% bovine serum albumin / TBS and then incubated with primary antibodies (rabbit anti α-actinin (Cell Signaling, 3134)) overnight at 4 °C at 1:30. The secondary antibody (donkey anti-rabbit-alexafluor-488 (Thermo Fisher, A-21206)), was incubated for 2h at room temperature at 1:500 together with alexa-coupled wheat germ agglutinin (WGA, 1:500, Thermofisher). Sections were mounted in DAPI Fluoromount-G (SouthernBiotech, USA).

Images were captured using a Leica TCS SP5 confocal microscope (Leica Microsystems) with ×40 HCX PL APO CS oil (NA= 1.3) objective. Three-dimensional images were collected, and a maximum projection image was created using the Leica LAS AF software.

### Statistical Analysis

Differences between experimental groups were analyzed using Microsoft Excel (v13.7) and GraphPad Prism (v9.1.1) statistical tools. Sample distribution was assumed normal with equal variance. Statistical analysis was performed using Student’s t-tests or one-way ANOVA with post-hoc Bonferroni-corrected t-tests and p<0.05 was considered to be statistically significant. Sample sizes of biologically independent samples per group and the number of independent experiments are indicated in Fig. legends.

## Author Contributions

DCA, YM designed the study with assistance from DR, CK, DM, JJ, DJ, HY, and RACL. DCA and YM designed the experiments. DCA supervised all the experiments, led the data analyses, and prepared the manuscript with YM. DR and CK performed the RNA Sequencing analyses. DL, HB, and DW performed COVID-19 autopsy harvesting and staining and provided feedback. DCA performed all the cell culture, contraction amplitude studies, ELISA, immunofluorescence staining, confocal imaging, and image analysis with assistance from CK and KT. NH created the schematics and assisted with the manuscript preparation. DM, DJ, JJ, RACL, and YM supervised the collective efforts of this study including manuscript preparation.

## Competing Interests

The authors declare no competing interests.

## Acknowledgements

The work is supported by the National Institutes of Health (R01HL133308, F31 HL154665), the National Science Foundation (EPS-0903795), the NIH Cardiovascular Training Grant (T32 HL007260), and US Department of Veterans Affairs Merit Review (I01 BX002327). This study used the services of the Morphology, Imaging, and Instrumentation Core, which is supported by NIH-NIGMS P30 GM103342 to the South Carolina COBRE for Developmentally Based Cardiovascular Diseases. This work was supported in part by the Translational Science Shared Resource, Hollings Cancer Center, Medical University of South Carolina (P30 CA138313).

## Notes

### Competing Interest Statement

The authors have declared no competing interest.

## References

1 Zheng, Y. Y., Ma, Y. T., Zhang, J. Y. & Xie, X. COVID-19 and the cardiovascular system. Nature reviews. Cardiology, doi:10.1038/s41569-020-0360-5 (2020).

2 Ruan, Q., Yang, K., Wang, W., Jiang, L. & Song, J. Clinical predictors of mortality due to COVID-19 based on an analysis of data of 150 patients from Wuhan, China. Intensive Care Med, doi:10.1007/s00134-020-05991-x (2020).

3 Zhou, F. et al. Clinical course and risk factors for mortality of adult inpatients with COVID-19 in Wuhan, China: a retrospective cohort study. The Lancet 395, 1054–1062, doi:10.1016/s0140-6736(20)30566-3 (2020).

4 Szekely, Y. et al. Spectrum of Cardiac Manifestations in COVID-19: A Systematic Echocardiographic Study. Circulation 142, 342–353, doi:10.1161/CIRCULATIONAHA.120.047971 (2020).

5 Lindner, D. et al. Association of Cardiac Infection With SARS-CoV-2 in Confirmed COVID-19 Autopsy Cases. JAMA Cardiol, doi:10.1001/jamacardio.2020.3551 (2020).

6 Juan A. Perez-Bermejo1†, S. K., Sarah J. Rockwood1†, Camille R. Simoneau1,2†, et al. SARS-CoV-2 infection of human iPSC–derived cardiac cells reflects cytopathic features in hearts of patients with COVID-19. Science Translational Medicine 13 (2021).

7 Zhu, H. et al. Cardiovascular Complications in Patients with COVID-19: Consequences of Viral Toxicities and Host Immune Response. Curr Cardiol Rep 22, 32, doi:10.1007/s11886-020-01292-3 (2020).

8 Bavishi, C. et al. Acute myocardial injury in patients hospitalized with COVID-19 infection: A review. Prog Cardiovasc Dis, doi:10.1016/j.pcad.2020.05.013 (2020).

9 Giustino, G. et al. Coronavirus and Cardiovascular Disease, Myocardial Injury, and Arrhythmia: JACC Focus Seminar. Journal of the American College of Cardiology 76, 2011–2023, doi:10.1016/j.jacc.2020.08.059 (2020).

10 Mehta, P. et al. COVID-19: consider cytokine storm syndromes and immunosuppression. The Lancet, doi:10.1016/s0140-6736(20)30628-0 (2020).

11 Murthy, H., Iqbal, M., Chavez, J. C. & Kharfan-Dabaja, M. A. Cytokine Release Syndrome: Current Perspectives. Immunotargets Ther 8, 43–52, doi:10.2147/ITT.S202015 (2019).

12 Fajgenbaum, D. C. & June, C. H. Cytokine Storm. The New England journal of medicine 383, 2255–2273, doi:10.1056/NEJMra2026131 (2020).

13 Brodin, P. Immune determinants of COVID-19 disease presentation and severity. Nature medicine 27, 28–33, doi:10.1038/s41591-020-01202-8 (2021).

14 Jenner, A. L. et al. COVID-19 virtual patient cohort reveals immune mechanisms driving disease outcomes. bioRxiv, doi:10.1101/2021.01.05.425420 (2021).

15 Lucas, C. et al. Longitudinal analyses reveal immunological misfiring in severe COVID-19. Nature 584, 463–469, doi:10.1038/s41586-020-2588-y (2020).

16 Moreno-Perez, O. et al. Experience with tocilizumab in severe COVID-19 pneumonia after 80 days of follow-up: A retrospective cohort study. J Autoimmun, 102523, doi:10.1016/j.jaut.2020.102523 (2020).

17 Cavalli, G. et al. Interleukin-1 blockade with high-dose anakinra in patients with COVID-19, acute respiratory distress syndrome, and hyperinflammation: a retrospective cohort study. The Lancet Rheumatology 2, e325–e331, doi:10.1016/s2665-9913(20)30127-2 (2020).

18 Huet, T. et al. Anakinra for severe forms of COVID-19: a cohort study. The Lancet Rheumatology 2, e393–e400, doi:10.1016/s2665-9913(20)30164-8 (2020).

19 Kalil, A. C. et al. Baricitinib plus Remdesivir for Hospitalized Adults with Covid-19. New England Journal of Medicine, doi:10.1056/NEJMoa2031994 (2020).

20 Stone, J. H. et al. Efficacy of Tocilizumab in Patients Hospitalized with Covid-19. The New England journal of medicine 383, 2333–2344, doi:10.1056/NEJMoa2028836 (2020).

21 Group, W. H. O. R. E. A. f. C.-T. W. et al. Association Between Administration of Systemic Corticosteroids and Mortality Among Critically Ill Patients With COVID-19: A Meta-analysis. JAMA 324, 1330–1341, doi:10.1001/jama.2020.17023 (2020).

22 De Luca, G. et al. GM-CSF blockade with mavrilimumab in severe COVID-19 pneumonia and systemic hyperinflammation: a single-centre, prospective cohort study. The Lancet Rheumatology 2, e465–e473, doi:10.1016/s2665-9913(20)30170-3 (2020).

23 Bronte, V. et al. Baricitinib restrains the immune dysregulation in patients with severe COVID-19. The Journal of clinical investigation 130, 6409–6416, doi:10.1172/JCI141772 (2020).

24 Aouba, A. et al. Targeting the inflammatory cascade with anakinra in moderate to severe COVID-19 pneumonia: case series. Ann Rheum Dis, doi:10.1136/annrheumdis-2020-217706 (2020).

25 Puntmann, V. O. et al. Outcomes of Cardiovascular Magnetic Resonance Imaging in Patients Recently Recovered From Coronavirus Disease 2019 (COVID-19). JAMA Cardiol, doi:10.1001/jamacardio.2020.3557 (2020).

26 Puntmann, V. & Nagel, E. Cardiac Involvement After Recovering From COVID-19-Reply. JAMA Cardiol, doi:10.1001/jamacardio.2020.5285 (2020).

27 Mitrani, R. D., Dabas, N. & Goldberger, J. J. COVID-19 cardiac injury: Implications for long-term surveillance and outcomes in survivors. Heart Rhythm 17, 1984–1990, doi:10.1016/j.hrthm.2020.06.026 (2020).

28 Richards, D. J. et al. Inspiration from heart development: Biomimetic development of functional human cardiac organoids. Biomaterials 142, 112–123, doi:10.1016/j.biomaterials.2017.07.021 (2017).

29 Richards, D. J. et al. Human cardiac organoids for the modelling of myocardial infarction and drug cardiotoxicity. Nature Biomedical Engineering 4, 446–462, doi:10.1038/s41551-020-0539-4 (2020).

30 Delorey, T. M. et al. COVID-19 tissue atlases reveal SARS-CoV-2 pathology and cellular targets. Nature, doi:10.1038/s41586-021-03570-8 (2021).

31 Norelli, M. et al. Monocyte-derived IL-1 and IL-6 are differentially required for cytokine-release syndrome and neurotoxicity due to CAR T cells. Nature medicine 24, 739–748, doi:10.1038/s41591-018-0036-4 (2018).

32 Libby, P. Interleukin-1 Beta as a Target for Atherosclerosis Therapy. Journal of the American College of Cardiology 70, 2278–2289, doi:10.1016/j.jacc.2017.09.028 (2017).

33 Sandstedt, J. et al. Human cardiac fibroblasts isolated from patients with severe heart failure are immune-competent cells mediating an inflammatory response. Cytokine 113, 319–325, doi:10.1016/j.cyto.2018.09.021 (2019).

34 Mako, V. et al. Proinflammatory activation pattern of human umbilical vein endothelial cells induced by IL-1beta, TNF-alpha, and LPS. Cytometry A 77, 962–970, doi:10.1002/cyto.a.20952 (2010).

35 Aoyagi, T. & Matsui, T. The Cardiomyocyte as a Source of Cytokines in Cardiac Injury. J Cell Sci Ther 2012, doi:10.4172/2157-7013.s5-003 (2011).

36 Cappanera, S. et al. When Does the Cytokine Storm Begin in COVID-19 Patients? A Quick Score to Recognize It. J Clin Med 10, doi:10.3390/jcm10020297 (2021).

37 Sang, C. J., 3rd et al. Cardiac pathology in COVID-19: a single center autopsy experience. Cardiovasc Pathol 54, 107370, doi:10.1016/j.carpath.2021.107370 (2021).

38 Yang, L. et al. An Immuno-Cardiac Model for Macrophage-Mediated Inflammation in COVID-19 Hearts. Circulation research 129, 33–46, doi:10.1161/CIRCRESAHA.121.319060 (2021).

39 Roshdy, A., Zaher, S., Fayed, H. & Coghlan, J. G. COVID-19 and the Heart: A Systematic Review of Cardiac Autopsies. Front Cardiovasc Med 7, 626975, doi:10.3389/fcvm.2020.626975 (2020).

40 Italia, L. et al. COVID-19 and Heart Failure: From Epidemiology During the Pandemic to Myocardial Injury, Myocarditis, and Heart Failure Sequelae. Front Cardiovasc Med 8, 713560, doi:10.3389/fcvm.2021.713560 (2021).

41 Dai, D. F., Danoviz, M. E., Wiczer, B., Laflamme, M. A. & Tian, R. Mitochondrial Maturation in Human Pluripotent Stem Cell Derived Cardiomyocytes. Stem Cells Int 2017, 5153625, doi:10.1155/2017/5153625 (2017).

42 Aziz, M., Fatima, R. & Assaly, R. Elevated interleukin-6 and severe COVID-19: A meta-analysis. J Med Virol 92, 2283–2285, doi:10.1002/jmv.25948 (2020).

43 Soy, M. et al. Cytokine storm in COVID-19: pathogenesis and overview of anti-inflammatory agents used in treatment. Clin Rheumatol 39, 2085–2094, doi:10.1007/s10067-020-05190-5 (2020).

44 Tang, Y. et al. Cytokine Storm in COVID-19: The Current Evidence and Treatment Strategies. Front Immunol 11, 1708, doi:10.3389/fimmu.2020.01708 (2020).

45 Liau, B., Christoforou, N., Leong, K. W. & Bursac, N. Pluripotent stem cell-derived cardiac tissue patch with advanced structure and function. Biomaterials 32, 9180–9187, doi:10.1016/j.biomaterials.2011.08.050 (2011).

46 Litvinukova, M. et al. Cells of the adult human heart. Nature 588, 466–472, doi:10.1038/s41586-020-2797-4 (2020).

47 Del Valle, D. M. et al. An inflammatory cytokine signature predicts COVID-19 severity and survival. Nature medicine 26, 1636–1643, doi:10.1038/s41591-020-1051-9 (2020).

48 Jin, Y. et al. Endothelial activation and dysfunction in COVID-19: from basic mechanisms to potential therapeutic approaches. Signal Transduct Target Ther 5, 293, doi:10.1038/s41392-020-00454-7 (2020).

49 Varga, Z. et al. Endothelial cell infection and endotheliitis in COVID-19. The Lancet 395, 1417–1418, doi:10.1016/s0140-6736(20)30937-5 (2020).

50 Escher, R., Breakey, N. & Lammle, B. Severe COVID-19 infection associated with endothelial activation. Thromb Res 190, 62, doi:10.1016/j.thromres.2020.04.014 (2020).

51 Juni, R. P. et al. Cardiac Microvascular Endothelial Enhancement of Cardiomyocyte Function Is Impaired by Inflammation and Restored by Empagliflozin. JACC Basic Transl Sci 4, 575–591, doi:10.1016/j.jacbts.2019.04.003 (2019).

52 Kyriazopoulou, E. et al. Early treatment of COVID-19 with anakinra guided by soluble urokinase plasminogen receptor plasma levels: a double-blind, randomized controlled phase 3 trial. Nat Med 27, 1752–1760, doi:10.1038/s41591-021-01499-z (2021).

53 Brauninger, H. et al. Cardiac SARS-CoV-2 infection is associated with pro-inflammatory transcriptomic alterations within the heart. Cardiovascular research, doi:10.1093/cvr/cvab322 (2021).

54 Ahmed, M. H. & Hassan, A. Dexamethasone for the Treatment of Coronavirus Disease (COVID-19): a Review. SN Compr Clin Med, 1–10, doi:10.1007/s42399-020-00610-8 (2020).

55 Group, R. C. et al. Dexamethasone in Hospitalized Patients with Covid-19. The New England journal of medicine 384, 693–704, doi:10.1056/NEJMoa2021436 (2021).

56 Tomazini, B. M. et al. Effect of Dexamethasone on Days Alive and Ventilator-Free in Patients With Moderate or Severe Acute Respiratory Distress Syndrome and COVID-19: The CoDEX Randomized Clinical Trial. JAMA 324, 1307–1316, doi:10.1001/jama.2020.17021 (2020).

57 Furlow, B. COVACTA trial raises questions about tocilizumab’s benefit in COVID-19. The Lancet Rheumatology 2, doi:10.1016/s2665-9913(20)30313-1 (2020).

58 Mills, R. J. et al. BET inhibition blocks inflammation-induced cardiac dysfunction and SARS-CoV-2 infection. Cell 184, 2167–2182 e2122, doi:10.1016/j.cell.2021.03.026 (2021).

59 Brotman, D. J. et al. Effects of short-term glucocorticoids on hemostatic factors in healthy volunteers. Thromb Res 118, 247–252, doi:10.1016/j.thromres.2005.06.006 (2006).

60 G. Sakuntala Warshamana, S. M. & Joseph A. Lasky, M. C., AND Arnold R. Brody. Dexamethasone activates expression of the PDGF-? receptor and induces lung fibroblast proliferation. American Physiological Society (1998).

61 Nalbandian, A. et al. Post-acute COVID-19 syndrome. Nature medicine, doi:10.1038/s41591-021-01283-z (2021).

62 Hartmann, C. et al. The Pathogenesis of COVID-19 Myocardial Injury: An Immunohistochemical Study of Postmortem Biopsies. Front Immunol 12, 748417, doi:10.3389/fimmu.2021.748417 (2021).

63 Huang, L. et al. Cardiac Involvement in Patients Recovered From COVID-2019 Identified Using Magnetic Resonance Imaging. JACC Cardiovasc Imaging 13, 2330–2339, doi:10.1016/j.jcmg.2020.05.004 (2020).

64 Lampropoulos, C. E. et al. Myocardial fibrosis after COVID-19 infection and severe sinus arrest episodes in an asymptomatic patient with mild sleep apnea syndrome: A case report and review of the literature. Respir Med Case Rep 32, 101366, doi:10.1016/j.rmcr.2021.101366 (2021).

65 Metzl, J. D. et al. Considerations for Return to Exercise Following Mild-to-Moderate COVID-19 in the Recreational Athlete. HSS J, 1–6, doi:10.1007/s11420-020-09777-1 (2020).

66 Anand Kumar, V. T., Linda Dee, Jeanne Olson, Eugene Uretz, and Joseph E. Parrillo. Tumor Necrosis Factora alpha and Interleukin 1B Are Responsible for In Vitro Myocardial Cell Depression Induced by Human Septic Shock Serum. J Exp Med 183, 949–958 (1996).

67 Jun-ichi Oyama, H. S., Hidetoshi Momii, Xiao-shu Cheng, Naoto Fukuyama,* Yukinori Arai, Kensuke Egashira, Hiroe Nakazawa,* and Akira Takeshita. Role of Nitric Oxide and Peroxynitrite in the Cytokine-induced Sustained Myocardial Dysfunction in Dogs In Vivo. J Clin Invest 101, 2207–2214 (1997).

68 Colliva, A., Braga, L., Giacca, M. & Zacchigna, S. Endothelial cell-cardiomyocyte crosstalk in heart development and disease. J Physiol 598, 2923–2939, doi:10.1113/JP276758 (2020).

69 Lopes, R. D. et al. Therapeutic versus prophylactic anticoagulation for patients admitted to hospital with COVID-19 and elevated D-dimer concentration (ACTION): an openlabel, multicentre, randomised, controlled trial. The Lancet 397, 2253–2263, doi:10.1016/s0140-6736(21)01203-4 (2021).

70 Richards, D. J. et al. Human cardiac organoids for the modelling of myocardial infarction and drug cardiotoxicity. Nature Biomedical Engineering, doi:10.1038/s41551-020-0539-4 (2020).

71 Kerr, C. M., Richards, D., Menick, D. R., Deleon-Pennell, K. Y. & Mei, Y. Multicellular Human Cardiac Organoids Transcriptomically Model Distinct Tissue-Level Features of Adult Myocardium. Int J Mol Sci 22, doi:10.3390/ijms22168482 (2021).

72 Dobin, A. et al. STAR: ultrafast universal RNA-seq aligner. Bioinformatics 29, 15–21, doi:10.1093/bioinformatics/bts635 (2013).

73 Robert C Gentleman, V. J. C., Douglas M Bates, Ben Bolstad, Marcel Dettling, Sandrine Dudoit, Byron Ellis, Laurent Gautier, Yongchao Ge, Jeff Gentry, Kurt Hornik, Torsten Hothorn, & Wolfgang Huber, S. I., Rafael Irizarry, Friedrich Leisch, Cheng Li, Martin Maechler, Anthony J Rossini, Gunther Sawitzki, Colin Smith, Gordon Smyth, Luke Tierney, Jean YH Yang and Jianhua Zhang. Bioconductor: open software development for computational biology and bioinformatics. Genome Biology 5 (2004).

74 Zhou, Y. et al. Metascape provides a biologist-oriented resource for the analysis of systems-level datasets. Nature communications 10, 1523, doi:10.1038/s41467-019-09234-6 (2019).

75 Zhang, Y., Parmigiani, G. & Johnson, W. E. ComBat-seq: batch effect adjustment for RNA-seq count data. NAR Genom Bioinform 2, lqaa078, doi:10.1093/nargab/lqaa078 (2020).

76 Leek JT, J. W., Parker HS, Fertig EJ, Jaffe AE, Zhang Y, Storey JD, Torres LC (2021).

77 Liberzon, A. et al. Molecular signatures database (MSigDB) 3.0. Bioinformatics 27, 1739–1740, doi:10.1093/bioinformatics/btr260 (2011).

78 Subramanian, A. et al. Gene set enrichment analysis: a knowledge-based approach for interpreting genome-wide expression profiles. Proceedings of the National Academy of Sciences of the United States of America 102, 15545–15550, doi:10.1073/pnas.0506580102 (2005).

79 Caplan, A. I. All MSCs are pericytes? Cell Stem Cell 3, 229–230, doi:10.1016/j.stem.2008.08.008 (2008).

80 Caplan, A. I. New MSC: MSCs as pericytes are Sentinels and gatekeepers. J Orthop Res 35, 1151–1159, doi:10.1002/jor.23560 (2017).

