## Supplemental Data for "Human Cardiac Organoids to Model COVID-19 Cytokine Storm Induced Cardiac Injuries"

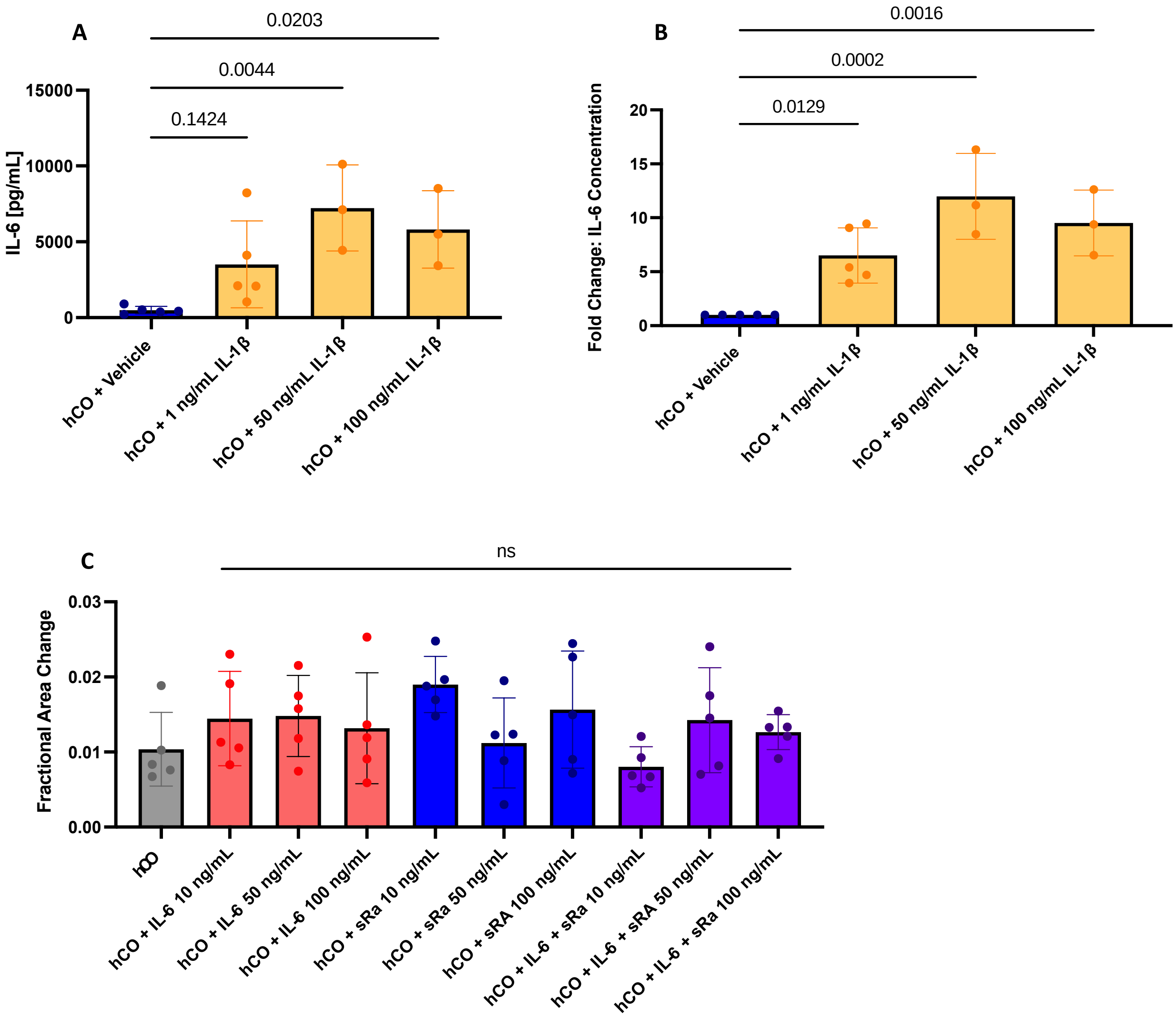


**Supplementary Figure 1. IL-6 is Present and Does Not Independently Alter Contraction Amplitude.** (A) Enzyme Linked Immunosorbent Assay (ELISA) indicating the release of IL-6 from hCOs in response to either 1 ng/mL, 50 ng/mL or 100 ng/mL of IL-1β. Fold Change is shown as mean +/- s.d., (n = 3-5). (B) ELISA results showing the concentration of IL-6 released in response to the same does of IL-1β in A. (C) FACs of hCOs on Day 10 after treatment with either IL-6 alone, the soluble IL-6 receptor (sRα), or the combination of IL-6 and sRα at 10 ng/mL, 50 ng/mL or 100 ng/mL. (n = 5, mean +/- s.d.). Statistical Analysis for each graph in this figure was performed by ANOVA.

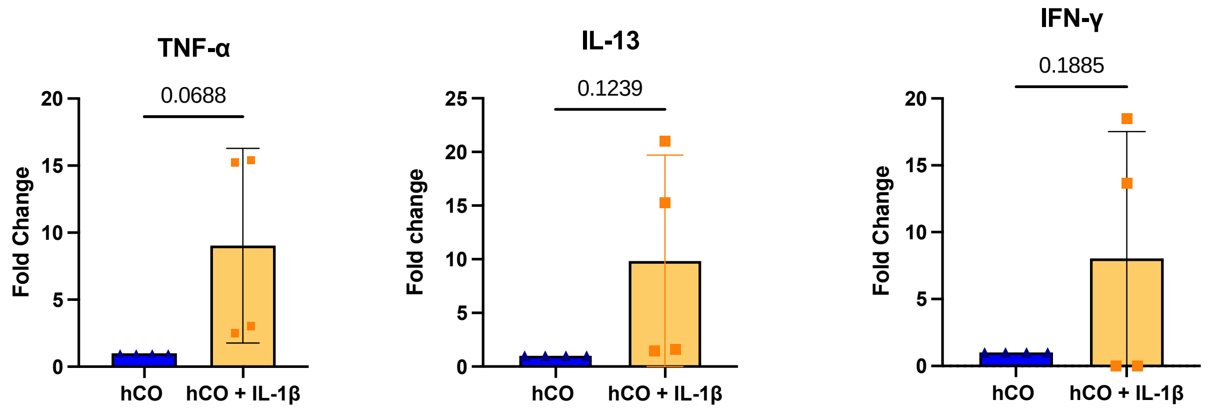


**Supplementary Figure 2. Additional Cytokines From Cytokine Multiplex Assay.** Fold change of cytokines in the hCO supernatant upon IL-1β treatment, relative to controls. (mean +/- s.d., n =4 for each group), analysis performed by student's t-test.

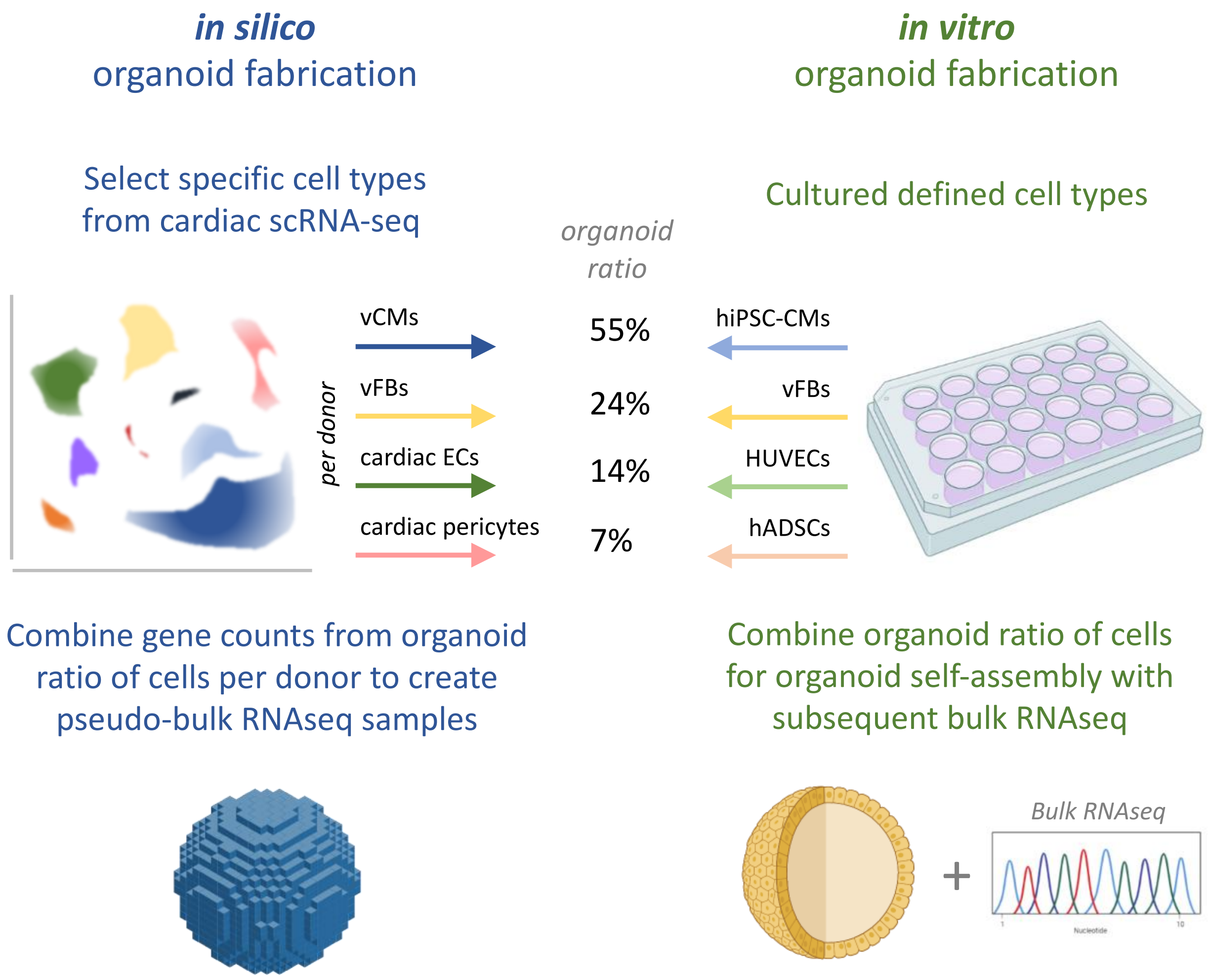


**Supplementary Figure 3: Schematic of *in silico* Organoid Fabrication**

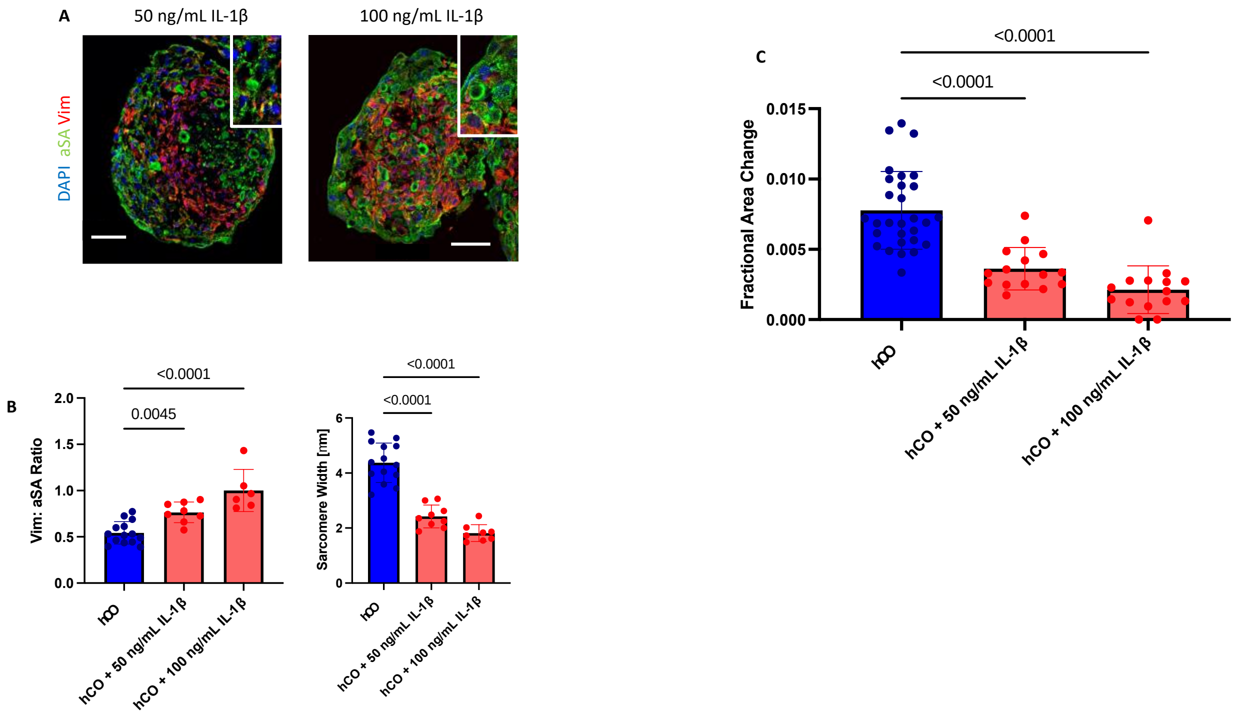


**Supplementary Figure 4. High Doses of IL-1β Show Similar Effects on Cardiomyocyte and Fibroblast Content as 1 ng/mL of IL-1β in Human Cardiac Organoids**. (A) Immunofluorescent staining of organoids at D4 after treatment with 50 ng/mL or 100 ng/mL of IL-1β. Top right indicates higher magnification image of their sarcomere structures. (Green = α-SA, Red = Vimentin, Blue = DAPI). (B) Quantification of immunofluorescent staining; (Left) Vimentin to α-SA ratio per organoid (n = 6-14), mean +/- s.d.), (Right) Mean sarcomere width in um for each treatment group. (n = 15-29, mean +/- s.d.). C. FAC on Day 4 of each treatment group (n= 15-30, mean +/- s.d.). Scale Bar for each image in figure = 50 um. Statistical Analysis performed by ANOVA for each group.

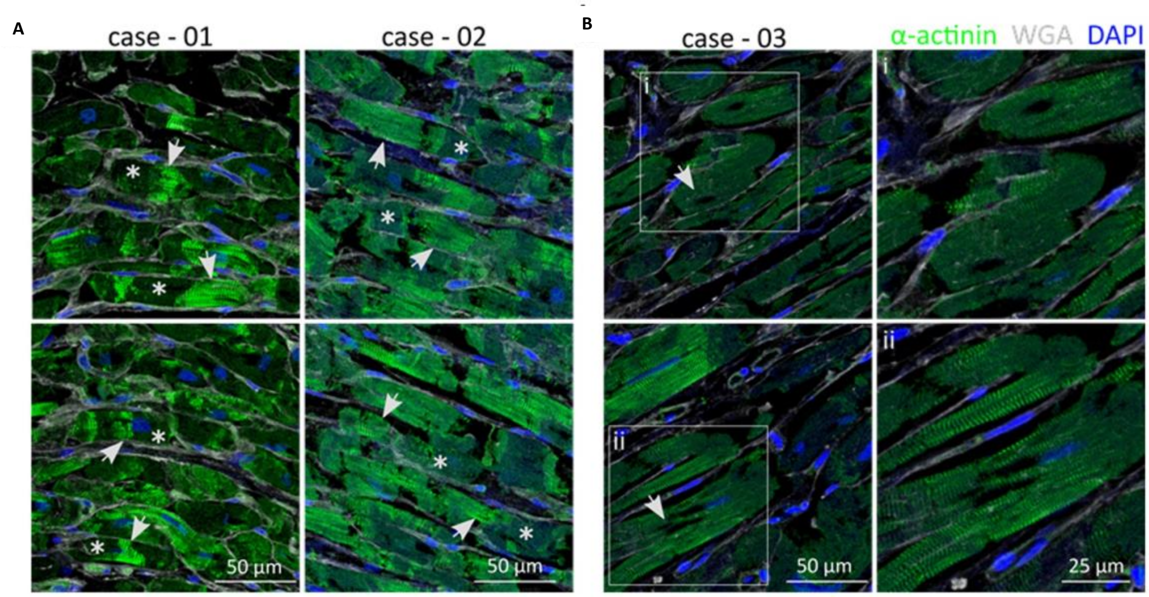


**Supplementary Figure 5: Histology of Hearts with Confirmed Viral Load**. (A) Two representative α-SA (green) of cardiac tissue section from case-01 and case-02 (both exhibiting high virus load of SARS-CoV-2 in the heart) are shown. The normal sarcomeric structure of cardiomyocytes is indicated by arrows, whereas inconsistent actinin staining as a sign of sarcomeric disarray is indicated by asterisks. Nuclei counterstained with DAPI are shown in blue, extracellular matrix stained with WGA is shown in white. (B) Two representative images of cardiac tissue from case-03 who exhibited high virus load in the heart. White boxes indicted zoomed in areas of the image on the right. Some cardiomyocytes lack nuclear DNA staining (DAPI). White arrows denote the putative region of nuclear localization.

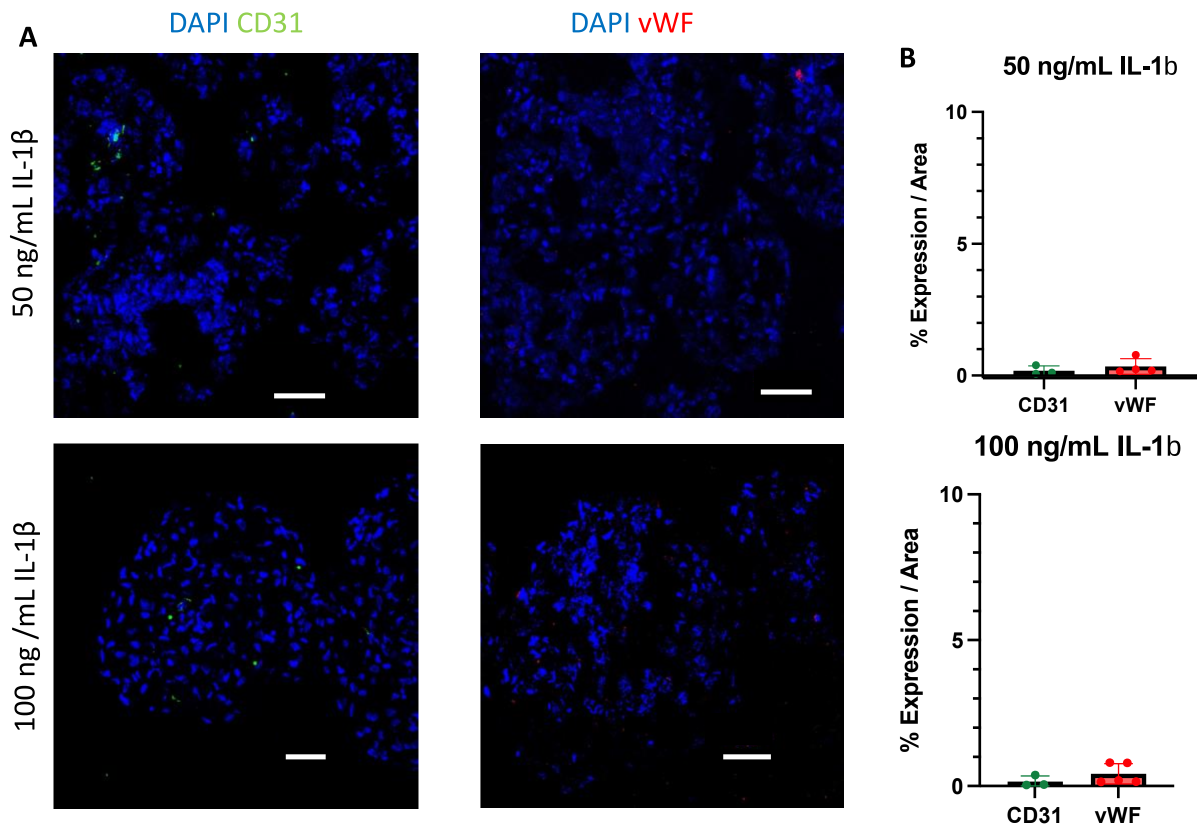


**Supplementary Figure 6: High Doses of IL-1β eliminate Endothelial Cell Content from Human Cardiac Organoids. (**A) Immunofluorescent Staining of hCOs treated with 50 ng/mL or 100 ng/mL of IL-1β on Day 4. (Green = CD31, Red = vWF, Blue = DAPI). (B) Quantification of CD31 and vwF expression for hCOs treated with 50 ng/mL or 100 ng/mL of IL-1β (n = 3-5, mean +/- s.d.).

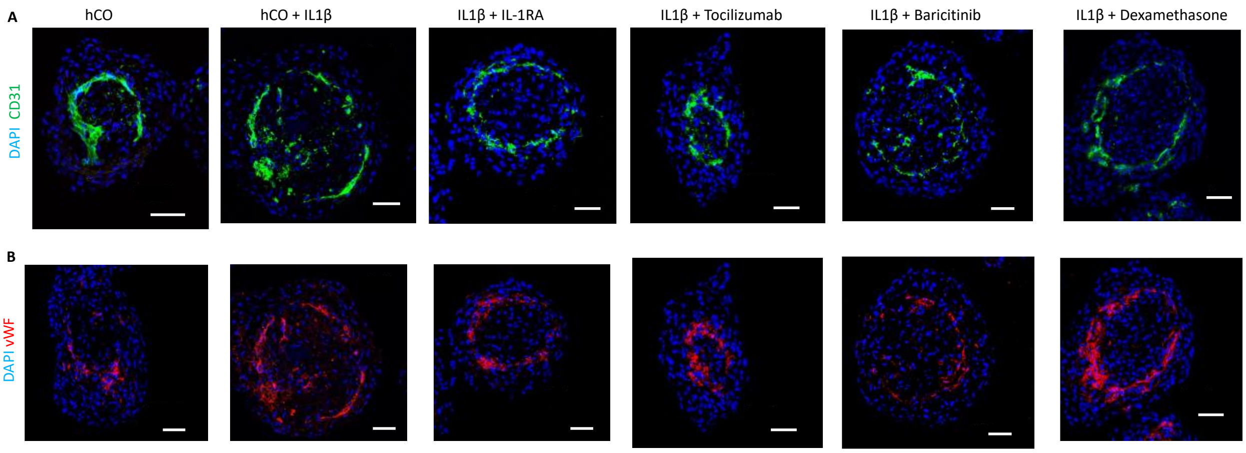


**Supplementary Figure 7. Representative Images of Vasculature Upon Treatment with Immunomodulatory Drugs.** (A) Immunofluorescent staining of hCOs on D4 with either vehicle, IL-1β, or IL-1β + each immunomodulatory drug (Green = CD31, Blue = DAPI). (B) Immunofluorescent staining of organoids on D4 with either vehicle, IL-1β, or IL-1β + each immunomodulatory drug (Red = vWF, Blue = DAPI). Scale bar for all images is 50 um.

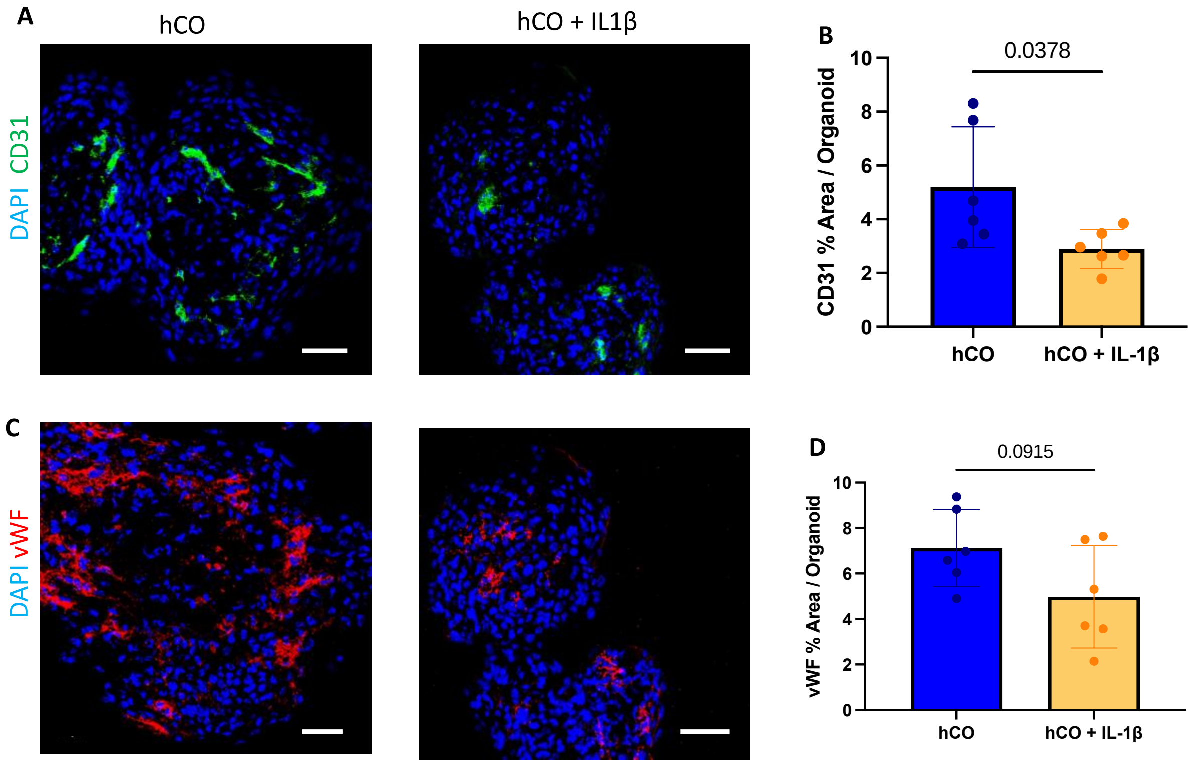


**Supplementary Figure 8: D14 Organoid Vasculature.** (A) Immunofluorescent Staining of hCOs after recovery culture on D14 (Green = CD31, Blue = DAPI). (B) Quantification of imaging seen in A. (C) Immunofluorescent Staining of hCOs after recovery culture on D14 (Red = vWF, Blue = DAPI). D. Quantification of staining seen in C. Scale bar for each image = 50 um. (n = 6, mean +/- s.d.), analysis performed by student's t-test.
